# TNAP and PHOSPHO1 function synergistically to afford critical control over the mineralisation of the postnatal murine skeleton

**DOI:** 10.64898/2026.02.25.707785

**Authors:** Lucie E Bourne, Aikta Sharma, Scott Dillon, Jacob Keen, Soher N Jayash, Natalie Crump, Lucinda AE Evans, Maya Karmali, Worachet Promruk, Claire E Clarkin, Sonoko Narisawa, Louise Stephen, Brian L Foster, José Luis Millán, Colin Farquharson, Katherine A Staines

**Affiliations:** Centre for Lifelong Health, School of Applied Sciences, University of Brighton, Brighton, BN2 4GT, UK; Department of Mechanical Engineering, University College London, London, WC1E 7JE, UK; Yusuf Hamied Department of Chemistry, University of Cambridge, Cambridge CB2 1EW, UK; School of Biological Sciences, University of Southampton, Southampton, SO17 1BJ, UK; The Roslin Institute and Royal (Dick) School of Veterinary Studies, University of Edinburgh, Easter Bush, Midlothian EH25 9RG, UK; Faculty of Life Sciences & Medicine, Kings College London, London, SE1 1UL, UK; Human Genetics Program, Sanford Burnham Prebys Medical Discovery Institute, La Jolla, CA 92037, USA; Division of Biosciences, College of Dentistry, The Ohio State University, Columbus, OH 43210, USA

**Keywords:** Tissue non-specific alkaline phosphatase, Phospho1, biomineralisation, bone, murine model

## Abstract

Biomineralisation is essential for skeletal integrity, yet the synergistic roles of tissue non-specific alkaline phosphatase (TNAP) and PHOSPHO1 in postnatal bone mineralisation remain poorly defined. To decipher this, we generated a novel murine model in which *Alpl* was deleted in *Prx1*- expressing cells (*Alpl*^*Prx1/Prx1*^) in mice with a global *Phospho1*^-/-^ deficiency to overcome the perinatal lethality that arises upon dual global deletion. Using a multi-modal approach to spatially phenotype the limbs of these animals, we reveal mice lacking both TNAP and PHOSPHO1 exhibit a distinct lack of mineralisation and altered anatomical structure at postnatal day 1 (PN1) and 3-weeks of age. Although viable, these mice did not thrive due to their reduced size, thus further investigations were conducted on mice with a heterozygous deletion of TNAP (*Alpl*^*wt/Prx1*^*;Phospho1*^*-/-*^). Although smaller than wild-types at PN1 and 3 weeks old, these mice did not display the gross limb deformations observed in the homozygous animals and the single, functioning *Alpl* allele rescued the loss of biomineralisation observed following dual phosphatase deletion. At 6-weeks of age, compromised epiphyses and metaphyses were only seen in *Alpl*^*Prx1/Prx1*^ animals. Further, we found that tibial geometry and porosity was significantly altered by *Phospho1* deletion (*Phospho1*^-/-^), which was compounded in the *Alpl*^*wt/Prx1*^*;Phospho1*^*-/-*^ mice and linked to alterations in collagen configuration, matrix mineralisation and growth plate deformities. Together, our findings establish the mechanistic framework for TNAP and PHOSPHO1 in permissive biomineralisation, providing critical insights into this fundamental process.

**Significance Statement:** Biomineralisation is essential for skeletal development and is critically dependent on phosphatases that release inorganic phosphate for hydroxyapatite formation. Our study investigates the dual role of PHOSPHO1 and TNAP in this process, using a novel murine knockout model. Deletion of both enzymes results in complete loss of bone mineralisation, demonstrating their critical synergistic function. Further, we show that PHOSPHO1 and TNAP exhibit distinct, spatially-restricted functions in the tibia and thus enhances our understanding of the fundamentals processes underpinning biomineralisation. These findings also have clinical relevance as they have the potential to inform on treatment strategies for hypo- and hyper-mineralised pathologies.

## Introduction

Biomineralisation of the skeleton is indispensable for vertebrate health and wellbeing throughout life. This involves a series of tightly regulated physicochemical and biochemical processes that facilitate the deposition of carbonated hydroxyapatite mineral crystals within the organic collagenous matrix of bone (1, 2). Local promoters of mineralisation are synthesised by osteoblasts and comprise of several phosphatases, with seminal studies showing tissue non-specific alkaline phosphatase (TNAP) and phosphoethanolamine/phosphocholine phosphatase 1 (PHOSPHO1) being central to this process (3-7). Although converging mechanisms between these phosphatases have been proposed (6, 8-10), how these interact and function to control the initiation and regulation of biomineralisation remains incomplete, representing a critical gap in our understanding of *de novo* bone formation.

Characterised by a hypomineralised bone matrix, missense mutations in the human TNAP (*ALPL*) gene lead to hypophosphatasia (HPP), a heritable form of rickets, and osteomalacia (4). Similarly, global deletion of *ALPL* in mice (*Alpl*^-/-^) phenocopies infantile hypophosphatasia. *Alpl*^-/-^ mice exhibit normal skeletal mineralisation at birth, yet by postnatal day 6, hypomineralisation becomes apparent and worsens until their early demise by postnatal day 20 (7). This aberrant mineralisation is, in part, driven by an accumulation of extracellular inorganic pyrophosphate (ePP_i_), a physiological substrate of TNAP and potent mineralisation inhibitor, which arrests mineral crystal propagation onto the collagen-rich extracellular matrix (ECM) (7, 11). ePP_i_ is produced ectoplasmically via cellular efflux of ATP from Ankylosis protein (ANK) and by the enzymatic action of ectonucleotide pyrophosphatase/phosphodiesterase-1 (ENPP1) that catabolises extracellular ATP into ePP_i_ and AMP (12-14). TNAP generates P_i_ from ePP_i_ while simultaneously restricting the concentration of ePP_i_ to maintain the PP_i_/P_i_ ratio permissive for bone mineralisation (12). TNAP can also promote matrix mineralisation by dephosphorylating the phosphorylated form of osteopontin (OPN), another potent mineralisation inhibitor (15).

To phenocopy late-onset HPP in skeletally mature animals, conditional deletion of *Alpl* using *Cre* recombinases in *Col1a1-* (osteoblasts and dental cells) and *Prx1-* (early limb-bud and craniofacial mesenchyme) expressing cells has revealed long bone skeletal defects, hypomineralisation and dentoalveolar disease (16). Small membrane bound matrix vesicles (MVs) provide an ideal environment for the formation of calcium-phosphate complexes, however, osteoblast-derived MVs in both hypophosphatasia patients and *Alpl*^-/-^ mice retain the ability to initiate intra-vesicular mineral formation; demonstrating that TNAP is not essential for the initiation of mineralisation (6, 17, 18).

PHOSPHO1 is the only known phosphatase specific to bone and cartilage (5, 19, 20). As a member of the haloacid dehalogenase superfamily of enzymes that functions as an intra-vesicular phosphatase, PHOSPHO1 exhibits high specificity towards phosphoethanolamine and phosphocholine (PCho) which are enriched in MV membranes (21-23). These PHOSPHO1 substrates enable the release of P_i_ inside MVs to promote mineral formation (22, 24). Genetic ablation of *Phospho1* in mice results in elevated ePP_i_ levels, osteomalacia, spontaneous fractures, spinal deformities and bowed long bones (6, 25, 26). In contrast to the skeletal phenotype of *Alpl* or *Phospho1*-deficient mice, the simultaneous global genetic ablation of both phosphatases results in an entirely unmineralised skeleton and perinatal death (6). This observation highlights that while TNAP and PHOSPHO1 function independently during matrix mineralisation, further interrogation of their roles will yield mechanistic insights into the initiation of skeletal mineralisation (6, 9).

Since mice lacking both *Alpl* and *Phospho1* die perinatally, it is unclear whether these phosphatases synergise to regulate bone mineralisation and skeletal structure in the postnatal skeleton. Mineralisation of the collagen matrix in the embryonic and postnatal skeleton differ in several key aspects and therefore need to be considered independently (27). Moreover, the growing postnatal skeleton is subject to extensive modelling events which is reliant on the mineralisation of newly formed osteoid. To fully comprehend the functional role of TNAP and PHOSPHO1 in the mineralisation of the postnatal skeleton, we circumvented the perinatal lethality of the *Alpl*^-/-^;*Phospho1*^-/-^ global knockout by generating mice with a conditional osteoblast-specific deletion of *Alpl* on a global *Phospho1*^-/-^ genetic background (*Alpl*^*Prx1/Prx1*^*;Phospho1*^*-/-*^).

## Results

### Dual deletion of *Alpl* and *Phospho1* results in complete lack of long bone mineralisation in postnatal male and female mice

Mice with a *Prx1*-specific deletion of TNAP were bred onto a global *Phospho1*^-/-^ background (*Alpl*^*Prx1/Prx1*^*;Phospho1*^*-/-*^). litters also consisted of wildtype (WT), *Prx1*-specific TNAP knockouts (*Alpl*^*Prx1/Prx1*^), *Phospho1* global knockouts (*Phospho1*^*-/-*^) and mice carrying a heterozygous *Prx1-* specific TNAP deletion on a *Phospho1*-null background (*Alpl*^*wt/Prx1*^*;Phospho1*^*-/-*^) (Fig. 1A).

**Figure 1.**
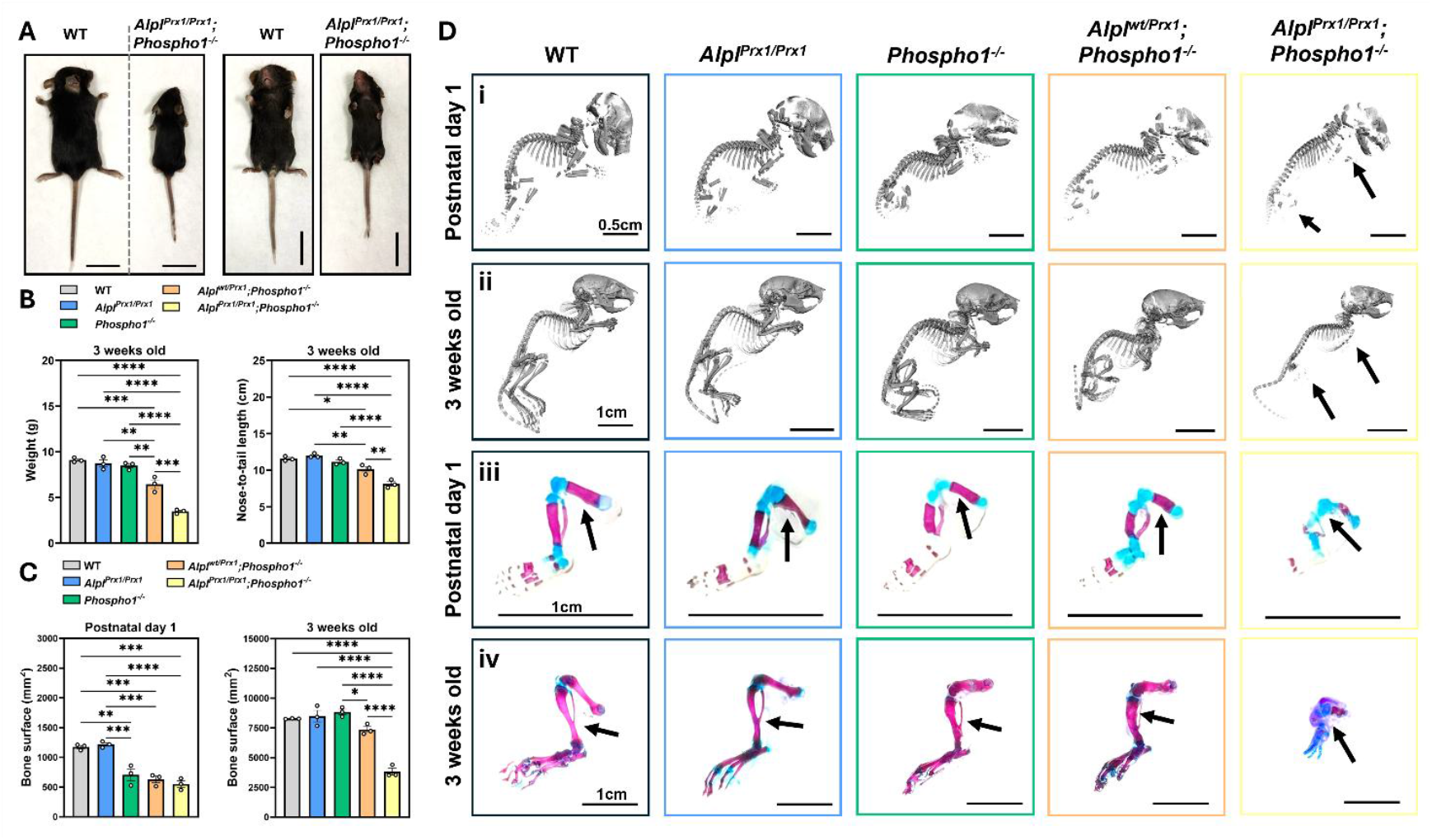
Gross phenotyping of mice with deletions in *Alpl* and *Phospho1* at postnatal day 1 and 3-weeks of age highlight severe malformations of the limbs and reduced size. (**A**) Schematic illustration of the genotypes generated for analysis, including mice that were wild-type (WT), had a *Prx1*-specific deletion of TNAP (*Alpl*^*Prx1/Prx1*^), a global deletion of PHOSPHO1 (*Phospho1*^*-/-*^), a heterozygous deletion of TNAP on this background (*Alpl*^*wt/Prx1*^*;Phospho1*^*-/-*^) and the double knockout (*Alpl*^*Prx1/Prx1*^*;Phospho1*^*-/-*^). (**B**) Back and front views of WT and *Alpl*^*Prx1/Prx1*^*;Phospho1*^*-/-*^ mice at 3-weeks of age. Scale bar = 2 cm. (**C**) Body weight and nose-to-tail lengths in 3-week-old mice. (**D**) Bone surface area of the whole skeleton analysed by µCT at postnatal day 1 and at 3-weeks old. (**E**) Representative µCT 3D-renderings of whole skeletons at (**i**) postnatal day 1 (scale bar = 5 mm) and (**ii**) 3 weeks old (scale bar = 1 cm). Arrows indicate lack of mineralised fore and hindlimbs in *Alpl*^*Prx1/Prx1*^*;Phospho1*^*-/-*^ mice. Whole mount imaging of hindlimbs in (**iii**) postnatal day 1 and (**iv**) 3 weeks old. Arrows indicate mineralisation in all genotypes except for *Alpl*^*Prx1/Prx1*^*;Phospho1*^*-/-*^ mice. Scale bar = 1 cm. Data are presented as mean ± SEM with points showing individual animals (n=3/genotype/age). *= *p*< 0.05, **= *p*< 0.01,***= *p*< 0.001, ****= *p*< 0.0001.

Western blotting indicated that the gene targeting strategy was successful with the expected absence of TNAP and/or PHOSPHO1 protein in the limbs of the various knockout mice (Fig. S1). *Alpl*^*Prx1/Prx1*^*;Phospho1*^*-/-*^ mice of both sexes were significantly smaller than their littermates, with severe limb and paw malformations evident at 3-weeks of age (Fig. 1B). Whilst *Alpl*^*wt/Prx1*^*;Phospho1*^*-/-*^ mice similarly weighed less and were smaller than WT, *Alpl*^*Prx1/Prx1*^ and *Phospho1*^*-/-*^ animals (*p*<0.05); *Alpl*^*Prx1/Prx1*^*;Phospho1*^*-/-*^ mice were significantly smaller than all other genotypes at 3-weeks of age (*p*<0.001; Fig. 1C).

Skeletal analysis by micro-computed tomography (µCT) at postnatal day 1 (PN1) showed that in comparison to WT and *Alpl*^*Prx1/Prx1*^ mice, deletion of *Phospho1* in all other genotypes resulted in reduced bone surface area (*p*<0.01; Fig. 1D).) The lack of mineralised limbs in *Alpl*^*Prx1/Prx1*^*;Phospho1*^*-/-*^ mice is evident in 3D-rendered images, whereas long bones and digits are observed in all other genotypes, but to a lesser extent in *Phospho1*^*-/-*^ and *Alpl*^*wt/Prx1*^*;Phospho1*^*-/-*^ mice (Fig. 1Ei). By 3-weeks of age, the mineralised surface area of *Phospho1*^*-/-*^ mice had recovered, whereas *Alpl*^*wt/Prx1*^*;Phospho1*^*-/-*^ mice maintained this reduction (*p*<0.05) and was further reduced in *Alpl*^*Prx1/Prx1*^*;Phospho1*^*-/-*^ mice (*p*<0.0001; Fig. 1D&Eii). Histological staining of skeletal elements from PN1 and 3-week-old mice with alcian blue and alizarin red confirmed a virtual absence of mineralised bone in *Alpl*^*Prx1/Prx1*^*;Phospho1*^*-/-*^ hindlimbs (Fig. 1Eiii&iv & Fig. S2). Visible limb malformations, including bowing and dysplasia, were observed in *Phospho1*^*-/-*^ animals, and to a greater extent in *Alpl*^*wt/Prx1*^*;Phospho1*^*-/-*^ mice (Fig. 1Eii&iv & Fig. S2).

Further histological analyses of hindlimbs from 3-week-old old mice revealed a lack of bone formation in the secondary ossification centre (SOC) of *Alpl*^*Prx1/Prx1*^, *Alpl*^*wt/Prx1*^*;Phospho1*^*-/-*^ and *Alpl*^*Prx1/Prx1*^*;Phospho1*^*-/-*^ mice, driven by *Alpl* loss (Fig. 2Ai-iii). Here, accumulations of hypertrophic chondrocytes surrounded by matrix persist, (Fig. 2Ai-iii, arrows) while the structure and composition of the articular cartilage appeared normal in all mice. Dual deletion of *Alpl* and *Phospho1* resulted in regionalised extension of the growth plate cartilage into the metaphyseal region (Fig. 2Ai&iii & Fig. S2C&D), containing SOX9- and MMP13-expressing chondrocytes (Fig. 2B&C). TRAP-positive osteoclasts were present at the osteochondral junction of all mice (Fig. 2D), but the metaphyseal tissue was devoid of clear trabeculae in *Alpl*^*Prx1/Prx1*^*;Phospho1*^*-/-*^ mice with instead, non-mineralised structures and minimal bone marrow evident (Fig. 2Ai-iii&D).

**Figure 2.**
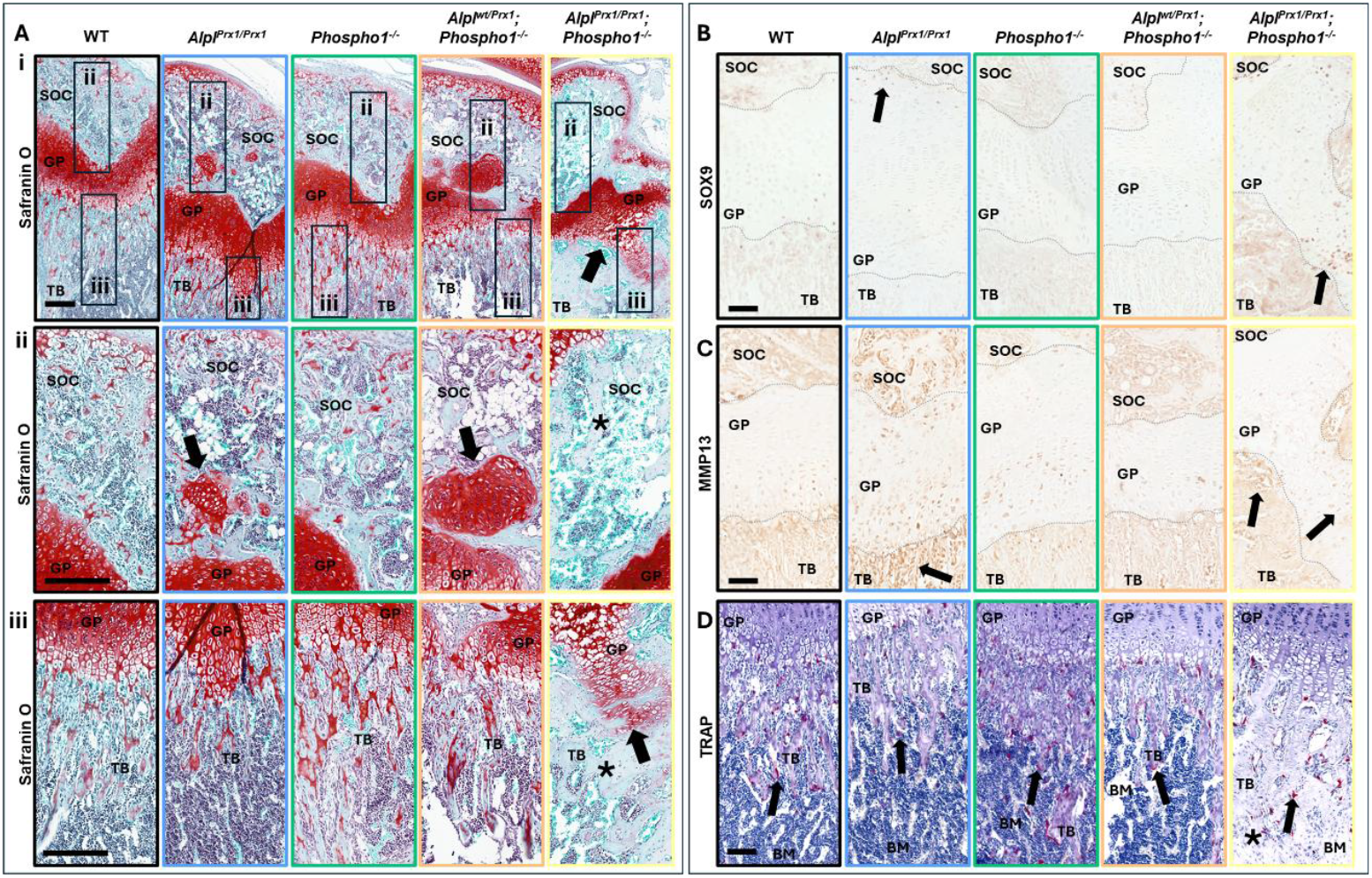
Histological analysis of 3-week-old mice with deletions in *Alpl* and *Phospho1* reveals abnormal bone formation in the secondary ossification centres, enlarged peripheral growth plate regions and a reduction in osteoclasts. (**Ai**) Representative images of Safranin-O/Fast green staining of femurs from wild-type (WT), *Alpl*^*Prx1/Prx1*^, *Phospho1*^*-/-*^, *Alpl*^*wt/Prx1*^*;Phospho1*^*-/-*^ and *Alpl*^*Prx1/Prx1*^*;Phospho1*^*-/-*^ mice. Images with a higher magnification of the SOC and TB are shown in Aii & iii. Arrows indicate accumulations of hypertrophic chondrocytes in the SOC and extension into the metaphysis. * indicates non-mineralised structures. Scale bar = 200 µm. (**B**) Representative SOX9 and (**C**) MMP13 immunolabelling. Arrows indicate disorganised chondrocytes in the GP and metaphyseal region. * indicates non-mineralised structures. The different regions are delineated by a dashed line. Scale bar = 100 µm. (**D**) Representative TRAP labelling of osteoclasts. Arrows indicate osteoclast expression in all animals, albeit reduced with *Alpl* and *Phospho1* deletion. Scale bar = 100 µm. GP = growth plate, SOC = secondary ossification centres, TB = metaphyseal trabecular bone.

### At skeletal maturity, one functional *Alpl* allele rescues biomineralisation and the epiphyseal defects observed with dual phosphatase deletion

Although viable, *Alpl*^*Prx1/Prx1*^*;Phospho1*^*-/-*^ mice did not thrive as well as their littermates post-weaning and remained stunted in their growth. In accordance with humane animal research practices in the UK, these mice were therefore not maintained beyond 3-weeks of age. Subsequent analyses were conducted on 6-week-old *Alpl*^*wt/Prx1*^*;Phospho1*^*-/-*^ heterozygous mice to allow delineation of the individual and dual roles of these phosphatases in the maturing skeleton.

Male and female mice of each genotype had comparable gross anatomy (Fig. S3A) and serum ALP and PP^i^ levels (Fig. S3B). However, while *Alpl*^*Prx1/Prx1*^ mice had similar body weight and lengths as WTs, *Phospho1*^*-/-*^ mice were smaller than WT mice (*p*<0.05; Table S1). *Alpl*^*wt/Prx1*^*;Phospho1*^*-/-*^ mice were also smaller in length than WTs, but body weight was comparable (Table S1). Analysis of bone lengths by µCT revealed that all genotypes (except *Alpl*^*Prx1/Prx1*^ males) had shorter tibias than WT mice, with progressively shorter bones in *Alpl*^*Prx1/Prx1*^ > *Phospho1*^*-/-*^ > *Alpl*^*wt/Prx1*^*;Phospho1*^*-/-*^ mice (*p*<0.05; Table S1).

The observed defects in the SOC of 3-week-old mice prompted further investigations of this region in 6-week-olds. Here, defects were only observed in *Alpl*^*Prx1/Prx1*^ mice. Morphometric analysis of the epiphyseal trabecular bone revealed reduced bone volume fraction (BV/TV), bone mineral density (BMD) and bone surface parameters in *Alpl*^*Prx1/Prx1*^ mice (*p*<0.0001, Fig. 3A-B & Fig. S4). Additional microarchitectural defects in the epiphysis including fewer trabeculae (Tb.N), increased spacing (Tb.Sp) and reduced complexity (Tb.Pf, Conn. Dn, Fractal dimension) were observed (*p*<0.001, Fig. 3A-B & Fig. S4). Von Kossa staining demonstrated that this reduction in mineralisation was accompanied by an increase in osteoid volume to bone volume (OV/BV) and thickness (O.Th) (*p*<0.01, Fig. 3C&D). Raman spectroscopy revealed alterations in epiphyseal ECM composition occur exclusively in female *Alpl*^*Prx1/Prx1*^ mice, as indicated by reductions in type I collagen content (sum peak area of proline and hydroxyproline) and enhanced collagen crosslinking, which was associated with low levels of B-type carbonate and modest reductions in hydroxyapatite (*p*<0.05, Fig. 3E & Fig. S5).

**Figure 3.**
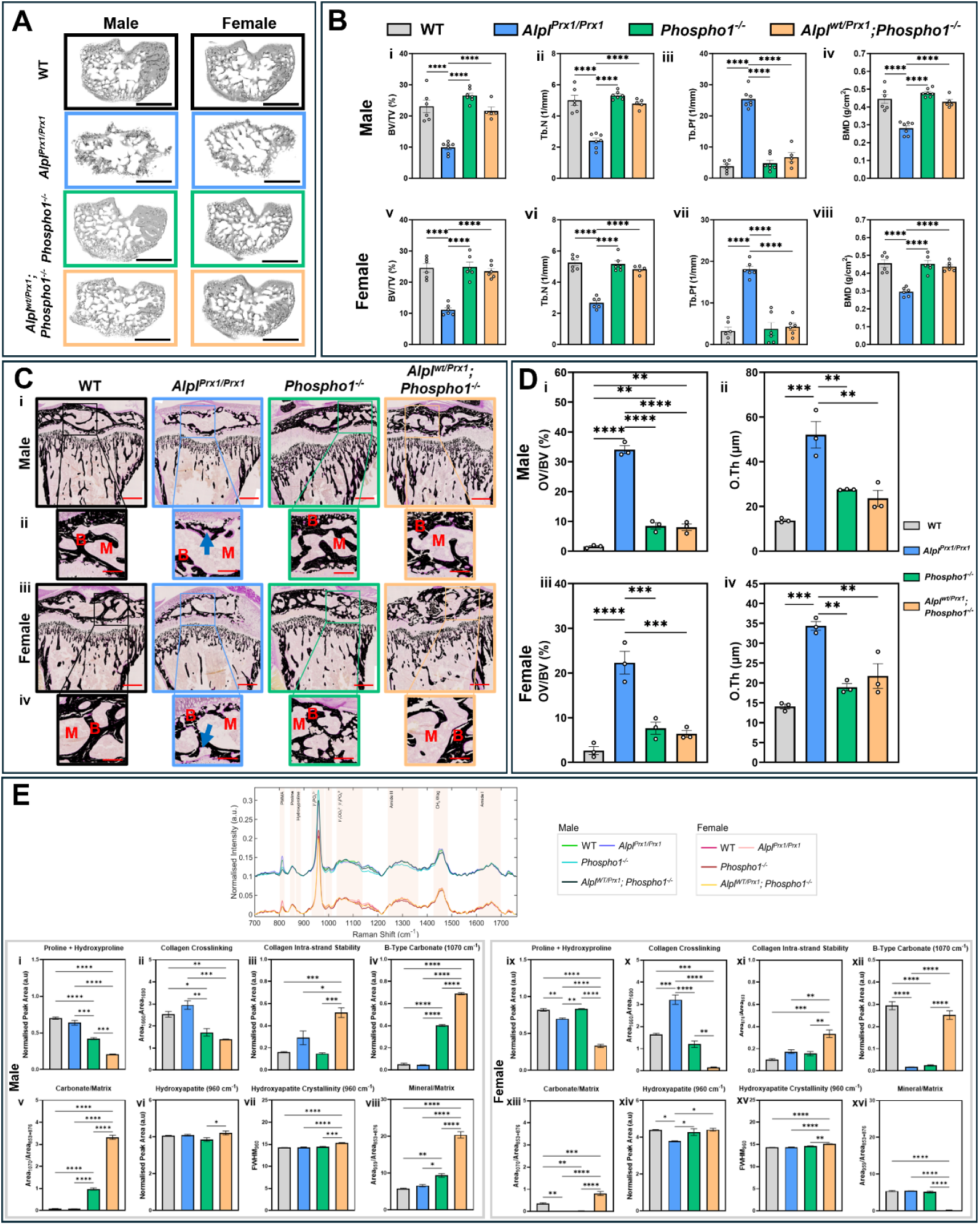
Tibial epiphyseal bone defects observed in 6-week-old mice with *Alpl* deletion and altered ECM composition in all genotypes. (**A**) Representative 3D-reconstructed μCT images of the epiphyseal trabecular region in wild-type (WT), *Alpl*^*Prx1/Prx1*^, *Phospho1*^*-/-*^ and *Alpl*^*wt/Prx1*^*;Phospho1*^*-/-*^ male and female mice. Scale bar = 1 mm. (**B**) Epiphyseal analysis of (**i & v**) bone volume fraction (BV/TV), (**ii & vi**) trabecular number (Tb.N), (**iii & vii**) trabecular pattern factor (Tb.Pf) and (**iv & viii**) bone mineral density (BMD) in males and females (n=5-7/genotype/sex). (**C**) Representative von Kossa-stained images of the epiphyseal region in (**i & ii**) male and (**iii & iv**) female mice. Arrows indicate areas of increased osteoid. B = bone, M = bone marrow. Scale bar = (**i & iii**) 500 µm; (**ii & iv**) 200 µm. (**D**) Quantification of (**i & iii**) osteoid volume/bone surface (OV/BV) and (**ii & iv**) osteoid thickness (O.Th) in the epiphyseal region of males and females, respectively (n=3/genotype/sex). Data are presented as mean ± SEM with points showing individual animals. (**E**) Deviations in vector normalised class means (n=75 spectra from 3 mice/genotype/sex) were detected in Raman bands corresponding to the ECM (proline, 853 cm^-1^; hydroxyproline, 876 cm^-1^; amide III region, 1242 cm^-1^; CH_2_ wag, 1450 cm^-1^ and amide I, 1660 cm^-1^) and mineral components (hydroxyapatite, 960 cm^-1^ and B-type carbonate, 1070 cm^-1^) of the tibial epiphyses. Spectral deconvolution of Raman bands in components of (**i & ix**) type I collagen (proline + hydroxyproline), (**ii & x**) collagen crosslinking, (**iii & xi**) collagen intra-strand stability, (**iv & xii**) B-type carbonate, (**v & xiii**) carbonate/matrix ratio, (**vi & xiv**) hydroxyapatite mineralisation and (**vii & xv**) crystallinity, and (**viii & xvi**) mineral/matrix ratio in males and females, respectively. Data are presented as the mean normalised peak area or full-width half maximum ± SEM. *= *p* < 0.05, **= *p* < 0.01, ***= *p* < 0.001, ****= *p* < 0.0001.

Further sex differences were evident in the ECM of *Phospho1*-deficient mice. Increased carbonation in both *Phospho1*^*-/-*^ and *Alpl*^*wt/Prx1*^*;Phospho1*^*-/-*^ males was accompanied by reduced type I collagen content and crosslinking (Fig. 3E & Fig. S5). In female *Phospho1*^*-/-*^ mice however, a sex-specific reduction in matrix carbonation occurred independently of changes to collagen and hydroxyapatite composition, whereas no change in carbonation was observed in female *Alpl*^*wt/Prx1*^*;Phospho1*^*-/-*^ animals, yet collagen crosslinking was impaired. Despite these differences, and in contrast to *Alpl*^*Prx1/Prx1*^ mice, *Phospho1* deletion had no effect on any epiphyseal morphometric parameters or von Kossa staining of mineralised tissue in either sex (Fig. 3A-D & Fig S4). Intriguingly, *Alpl*^*wt/Prx1*^*;Phospho1*^*-/-*^ mice showed no mineralisation abnormalities in the epiphysis (Fig. 3A-D & Fig S4), suggesting a potential correction of the observed histological differences at 3-weeks of age.

The metaphyseal trabecular regions of *Alpl*^*Prx1/Prx1*^ mice were also deleteriously affected, with reductions in BV/TV and BMD noted alongside disrupted trabecular microarchitecture (*p*<0.05; Fig. 4A-B & Fig S6). Osteoid mineralisation was also impaired in these animals (*p*<0.01; Fig. 4C&D), which may be driven by increased osteoclast number and associated bone (re)modelling (*p*<0.05; Fig. 4E-F). Whilst morphometric analyses in this region of *Phospho1*^*-/-*^ mice were unchanged (Fig. 4A-B), an increase in trabecular parameters was observed exclusively in female *Alpl*^*wt/Prx1*^*;Phospho1*^*-/-*^ mice (*p*<0.05; Fig. 4Bv-viii & Fig. S6). Despite conservation of osteoid levels, a trend towards reduced osteoclast number and surface was evident in male and female *Phospho1*^*-/-*^ and *Alpl*^*wt/Prx1*^*;Phospho1*^*-/-*^ mice (Fig. 4C-F).

**Figure 4.**
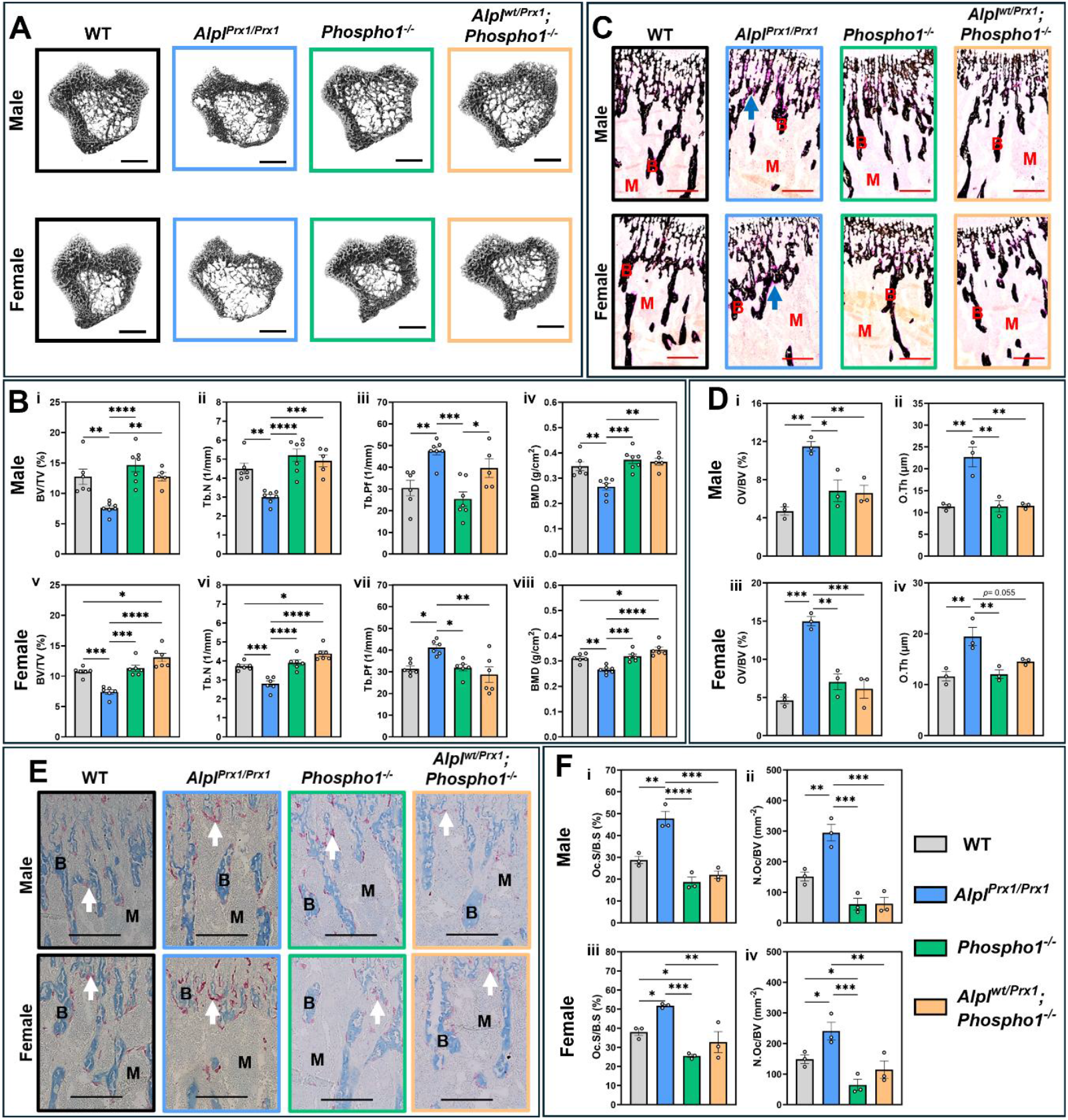
Tibial metaphyseal region analysis reveals trabecular compartment defects with *Alpl* deletion in 6-week-old mice, coupled with mineralisation defects and increased osteoclasts. (**A**) Representative 3D-reconstructed μCT images of the metaphyseal trabecular region in wild-type (WT), *Alpl*^*Prx1/Prx1*^, *Phospho1*^*-/-*^ and *Alpl*^*wt/Prx1*^*;Phospho1*^*-/-*^ male and female mice. Scale bar = 1 mm. (**B**) Trabecular analysis of (**i & v**) bone volume fraction (BV/TV), (**ii & vi**) trabecular number (Tb.N), (**iii & vii**) trabecular pattern factor (Tb.Pf) and (**iv & viii**) bone mineral density (BMD) in males and females, respectively (n=5-7/genotype/sex). (**C**) Representative von Kossa-stained images of the trabecular region. Arrows indicate areas of increased osteoid. B = bone, M = bone marrow. Scale bar = 200 µm. (**D**) Quantification of metaphyseal trabecular (**i & iii**) osteoid volume/bone volume (OV/BV) and (**ii & iv**) osteoid thickness (O.Th) in males and females, respectively (n=3/genotype/sex). (**E**) Representative images of TRAP labelling in the metaphyseal region. Arrows indicate red-labelled osteoclasts. B = bone, M = bone marrow. Scale bar = 200 µm. (**F**) Quantification of metaphyseal (**i & iii**) osteoclast surface/bone surface (Oc.S/B.S) and (**ii & iv**) number of osteoclasts/bone volume (N. OC/BV) in male and female mice, respectively (n=3/genotype/sex). Data are presented as mean ± SEM with points showing individual animals. *= *p* < 0.05, **= *p* < 0.01, ***= *p* < 0.001, ****= *p* < 0.0001.

3D reconstructions of *Alpl*^*Prx1/Prx1*^ growth plates further revealed malformation of the SOC (Fig. 5A & B). Here, thickness and volume of these reconstructions, alongside growth plate zone height, were unchanged, while elevations in growth plate bridge number in males was observed (Fig. 5B&C). This was accompanied by the accumulation of enlarged and disorganised chondrocytes in both the proliferative and hypertrophic regions (Fig. 5Di). In contrast, male *Phospho1*^*-/-*^ mice produced thicker growth plates, predominantly on the medial aspect of the condyle (Fig. 5A-C). This was linked to an expansion of the growth plate, where the formation of ‘pit-like’ regions containing disorganised, SOX9-expressing hypertrophic chondrocyte populations were evident (Fig. 5A&Dii). These trends were also observed in male *Alpl*^*wt/Prx1*^*;Phospho1*^*-/-*^ mice, yet the pit-like structures were not observed in females of either genotype.

**Figure 5.**
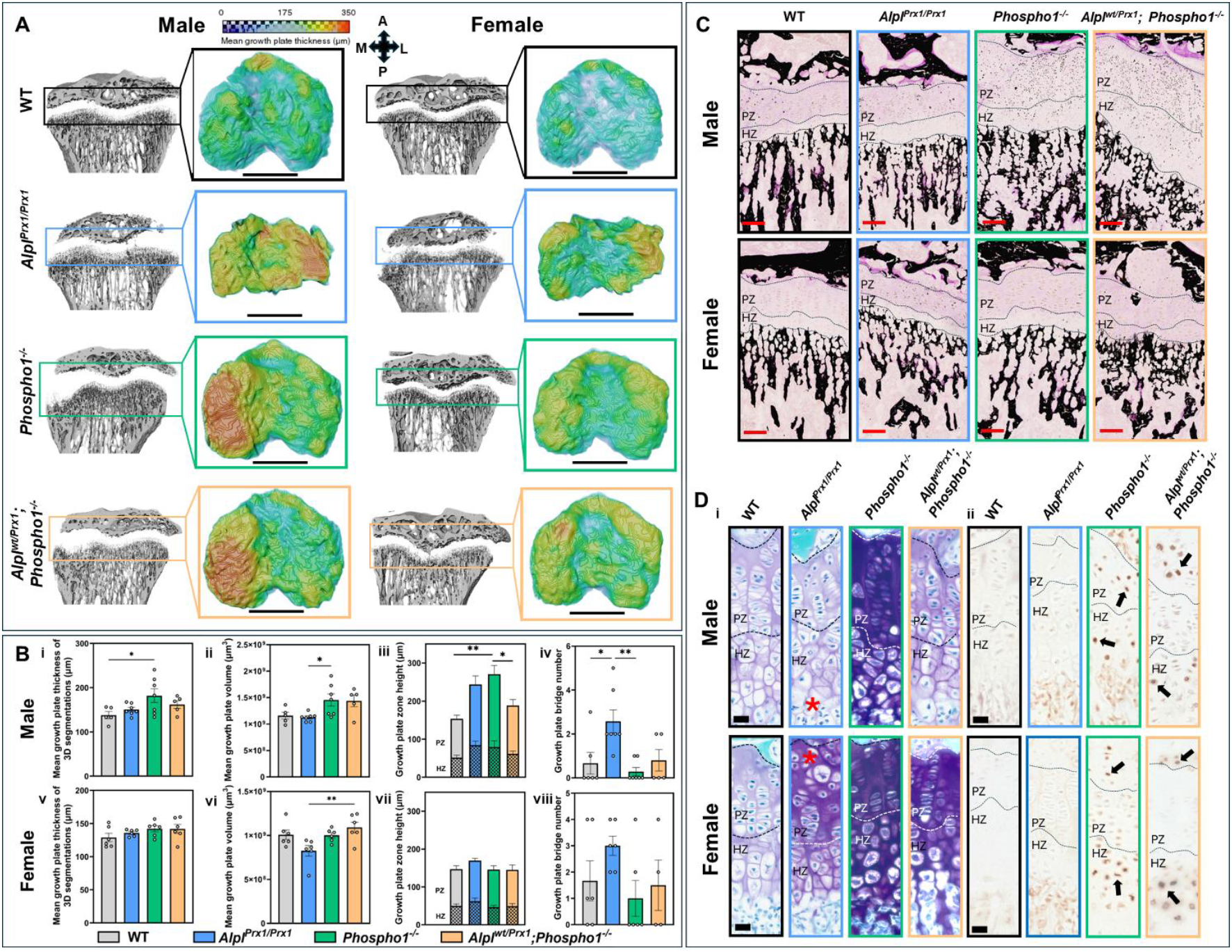
At 6 weeks of age, all genotypes display altered growth plate morphology. (**A**) Representative 3D-rendered images of the segmented region of the growth plate from wild-type (WT), *Alpl*^*Prx1/Prx1*^, *Phospho1*^*-/-*^ and *Alpl*^*wt/Prx1*^*;Phospho1*^*-/-*^ male and female mice. Scale bar = 1mm. (B) Measurements of the (**i & v**) mean growth plate thickness and (**ii & vi**) volume of the 3D segmentations in A, (**iii & vii**) growth plate zone height, and (**iv & viii**) number of bridges in males and females, respectively (n=5-7/genotype/sex). (**C**) Representative von Kossa-stained images of the growth plate. Proliferative (PZ) and hypertrophic (HZ) growth plate zones are delineated. Scale bar = 100μm. (**Di**) Representative Toluidine Blue-stained images of the growth plate and (**ii**) SOX9 immunolabelling in males and females. Proliferative (PZ) and hypertrophic (HZ) growth plate zones are delineated. * indicates enlarged chondrocytes. Arrows indicate SOX9 positive chondrocytes. Scale bar = 100μm. Data are presented as mean ± SEM with points showing individual animals. *= *p* < 0.05, **= *p* < 0.01.

### Deletion of *Phospho1* results in tibial geometry changes, which are compounded upon the additional deletion of one *Alpl* allele

To assess the effect of *Alpl* and *Phospho1* deletion along the full tibial length, 2D whole bone analysis and three-point bending were performed. Compared to WT mice, cortical bone area fraction (B.Ar/T.Ar) and thickness of male and female mice were reduced in all genotypes (Fig. 6A), leading to detrimental effects on the tibial mechanical properties in *Alpl*^*Prx1/Prx1*^ mice of both sexes, female *Phospho1*^*-/-*^, and female *Alpl*^*wt/Prx1*^*;Phospho1*^*-/-*^ animals (Table 1). Parameters associated with bone shape (predicted resistance to torsion (*J*), maximum and minimum moments of inertia (*I*_*max &*_ *I*_*min*_)), and tissue and bone perimeter (T.Pm and B.Pm, respectively) were increased in *Phospho1*^*-/-*^ mice compared to WT and *Alpl*^*Prx1/Prx1*^ (Fig. 6A & Fig. S7). In *Alpl*^*wt/Prx1*^*;Phospho1*^*-/-*^ animals, these effects were compounded, particularly at the most distal end and likely due to the severe bowing observed in this region (Fig. 6A&B). Whilst tissue mineral density (TMD) was also reduced in *Phospho1*^*-/-*^ and *Alpl*^*wt/Prx1*^*;Phospho1*^*-/-*^ mice (*p*<0.05; Fig. 6C), male mice of these genotypes were unaffected by mechanical testing (Table 1). Total porosity was increased along the tibial length in *Phospho1*^*-/-*^ and *Alpl*^*wt/Prx1*^*;Phospho1*^*-/-*^ male and female mice (Fig. 6Aiv&viii), whilst further analysis specifically at the tibiofibular junction identified increased canal and volume density (Fig. 6D).

**Table 1.**
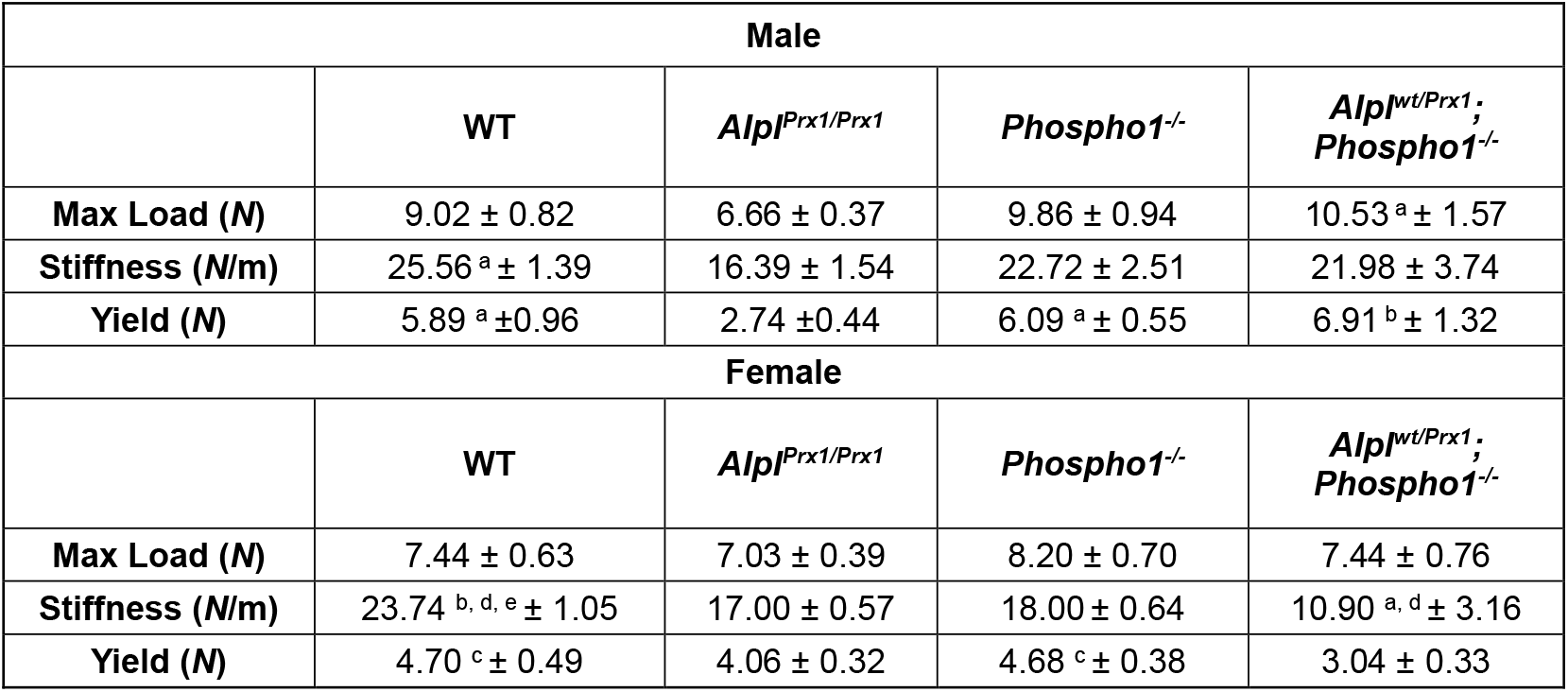
Three-point bending analysis of 6-week-old femurs from males and females. Data represented as mean ± SEM for *n*=5-7/genotype/sex. a = vs. *Alpl*^*Prx1/Prx1*^ (*), b = vs. *Alpl*^*Prx1/Prx1*^ (**), c = vs. *Alpl*^*wt/Prx1*^*;Phospho1*^*-/-*^ (*), d = vs. *Phospho1*^*-/-*^ (*), and e = vs. *Alpl*^*wt/Prx1*^*;Phospho1*^*-/-*^ (****).

| Male |  |  |  |  |
| --- | --- | --- | --- | --- |
|  | WT | *Alpl<sup>Prx1/Prx1</sup>* | *Phospho1<sup>-/-</sup>* | *Alpl<sup>wt/Prx1</sup>;Phospho1<sup>-/-</sup>* |
| Max Load (N) | 9.02 ± 0.82 | 6.66 ± 0.37 | 9.86 ± 0.94 | 10.53 <sup>a</sup> ± 1.57 |
| Stiffness (N/m) | 25.56 <sup>a</sup> ± 1.39 | 16.39 ± 1.54 | 22.72 ± 2.51 | 21.98 ± 3.74 |
| Yield (N) | 5.89 <sup>a</sup> ± 0.96 | 2.74 ± 0.44 | 6.09 <sup>a</sup> ± 0.55 | 6.91 <sup>b</sup> ± 1.32 |
| Female |  |  |  |  |
|  | WT | *Alpl<sup>Prx1/Prx1</sup>* | *Phospho1<sup>-/-</sup>* | *Alpl<sup>wt/Prx1</sup>;Phospho1<sup>-/-</sup>* |
| Max Load (N) | 7.44 ± 0.63 | 7.03 ± 0.39 | 8.20 ± 0.70 | 7.44 ± 0.76 |
| Stiffness (N/m) | 23.74 <sup>b, d, e</sup> ± 1.05 | 17.00 ± 0.57 | 18.00 ± 0.64 | 10.90 <sup>a, d</sup> ± 3.16 |
| Yield (N) | 4.70 <sup>c</sup> ± 0.49 | 4.06 ± 0.32 | 4.68 <sup>c</sup> ± 0.38 | 3.04 ± 0.33 |

**Figure 6.**
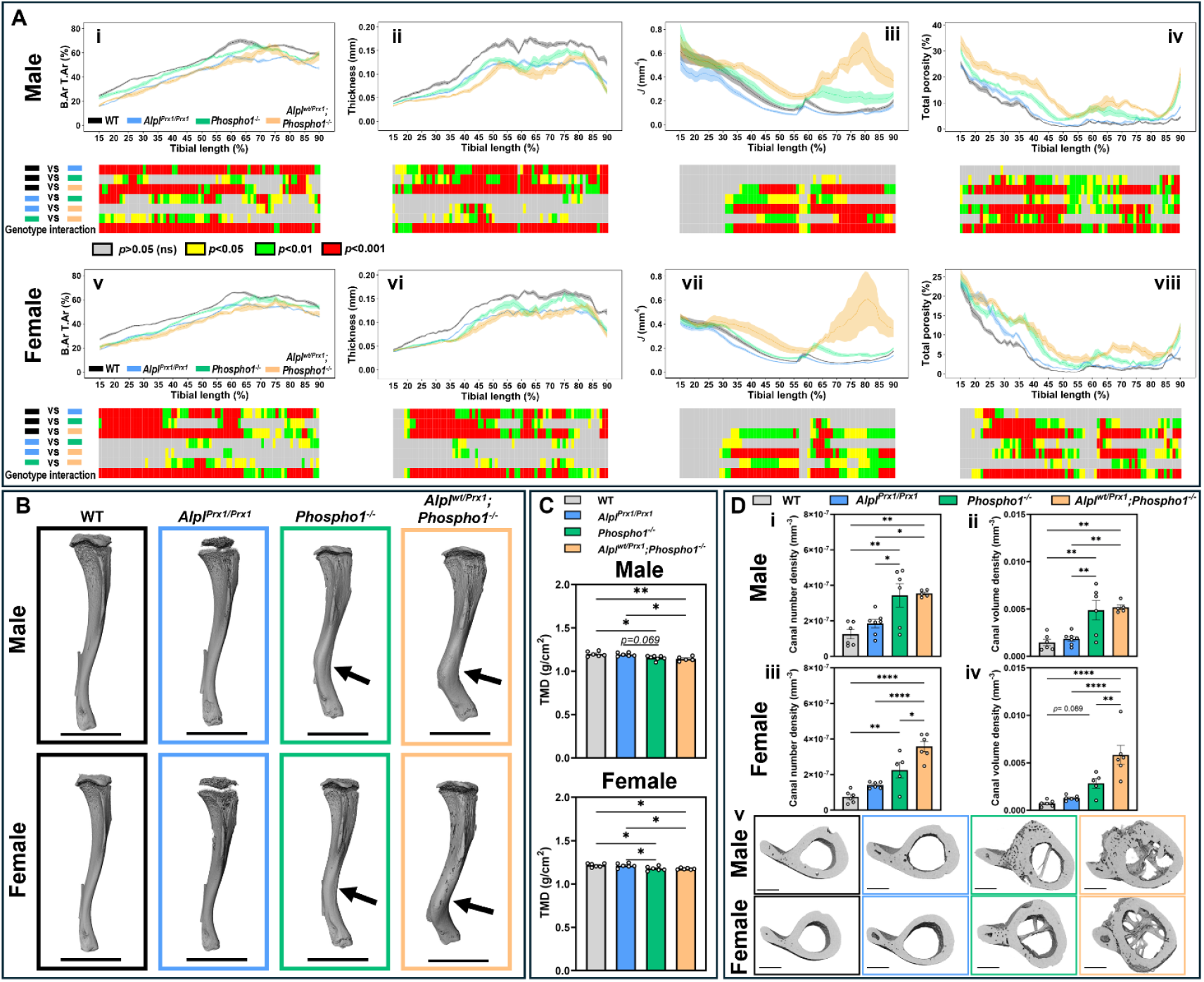
In 6-week-old mice, all genotypes exhibit reduced tibial cortical bone area and thickness, but *Phospho1* deficient animals have severely altered geometry and a mineralisation defect. (**A**) Measurement and statistical analysis heat map for (**i & v**) bone area / tissue area (B. Ar T. Ar), (**ii & vi**) thickness, (**iii & vii**) predicted resistance to torsion (*J*), and (**iv & viii**) total porosity in male and female mice, respectively. Line graphs represent mean ± SEM for wild-type (WT; black), *Alpl*^*Prx1/Prx1*^ (blue), *Phospho1*^*-/-*^ (green) and *Alpl*^*wt/Prx1*^*;Phospho1*^*-/-*^ (orange) male and female mice (n=5-7/genotype/sex). Graphical heat map summarises statistical differences at specific matched locations along the tibial length (15–90%). Red = *p* < 0.001, green = *p* < 0.01, yellow = *p* < 0.05, grey = *p* > 0.05 (not significant). (**B**) Representative 3D-rendered μCT images of the entire tibia. Arrows indicate bowing of the limb. Scale bar = 0.5 cm. (C) Quantification of tissue mineral density (TMD, g/cm^2^) in male and female mice. (**D**) Quantification of (**i&iii**) canal density and (**ii&iv**) volume density at the tibiofibular junction in males and females, respectively. (**v**) Representative 3D-reconstructed μCT images in males and females showing the porosity at the tibiofibular junction. Scale bar = 500 µm. Data are presented as mean ± SEM with points showing individual animals (n=5-7/genotype/sex). *= *p* < 0.05, **= *p* < 0.01.

To assess the mechanisms underpinning these alterations in cortical architecture and geometry, the expression of key genes involved in osteoblast differentiation and mineralisation were investigated in femoral diaphyseal bone. Interestingly, expression of *Enpp1, Bglap* (Osteocalcin) and *Spp1* (OPN) were downregulated in *Alpl*^*Prx1/Prx1*^ femurs compared to WT (*p*<0.05; Fig. 7A). Due to the nonsense mutation at amino acid 74 in exon 3 of *Phospho1*^*-/-*^ mice, gene primers for *Phospho1* expression could only be used on *Alpl*^*Prx1/Prx1*^ animals, which showed a reduction in *Phospho1* expression in both sexes (*p*<0.05; Fig. S8A). Genes associated with membrane PCho synthesis (*Chkb, Smpd3*) and phosphate transport (*Slc20a1, Slc20a2*) were generally elevated in *Phospho1*^*-/-*^ mice, whilst *Ank* and *Phex* were only increased in *Phospho1*^*-/-*^ and *Alpl*^*wt/Prx1*^*;Phospho1*^*-/-*^ males, respectively (*p*<0.05; Fig. S8A). Despite gene expression changes being predominantly affected by *Alpl*-loss, von Kossa staining of the cortical region revealed impaired mineralisation and an increase in osteoid in *Phospho1*^*-/-*^ and *Alpl*^*wt/Prx1*^*;Phospho1*^*-/-*^ male and female mice (Fig. 7B&C). The presence of osteoid was most obvious in *Alpl*^*wt/Prx1*^*;Phospho1*^*-/-*^ animals showing large areas of non-mineralised cortical tissue. Back scattered scanning electron microscopy (BSSEM) of the cortices confirmed this aberrant mineralisation in *Phospho1*^*-/-*^ and *Alpl*^*wt/Prx1*^*;Phospho1*^*-/-*^mice, with these animals displaying hypomineralised regions, large voids and inconsistent patchy regions of osteoid, highlighting a clear mineralisation defect (Fig. 7D & Fig. S8B).

**Figure 7.**
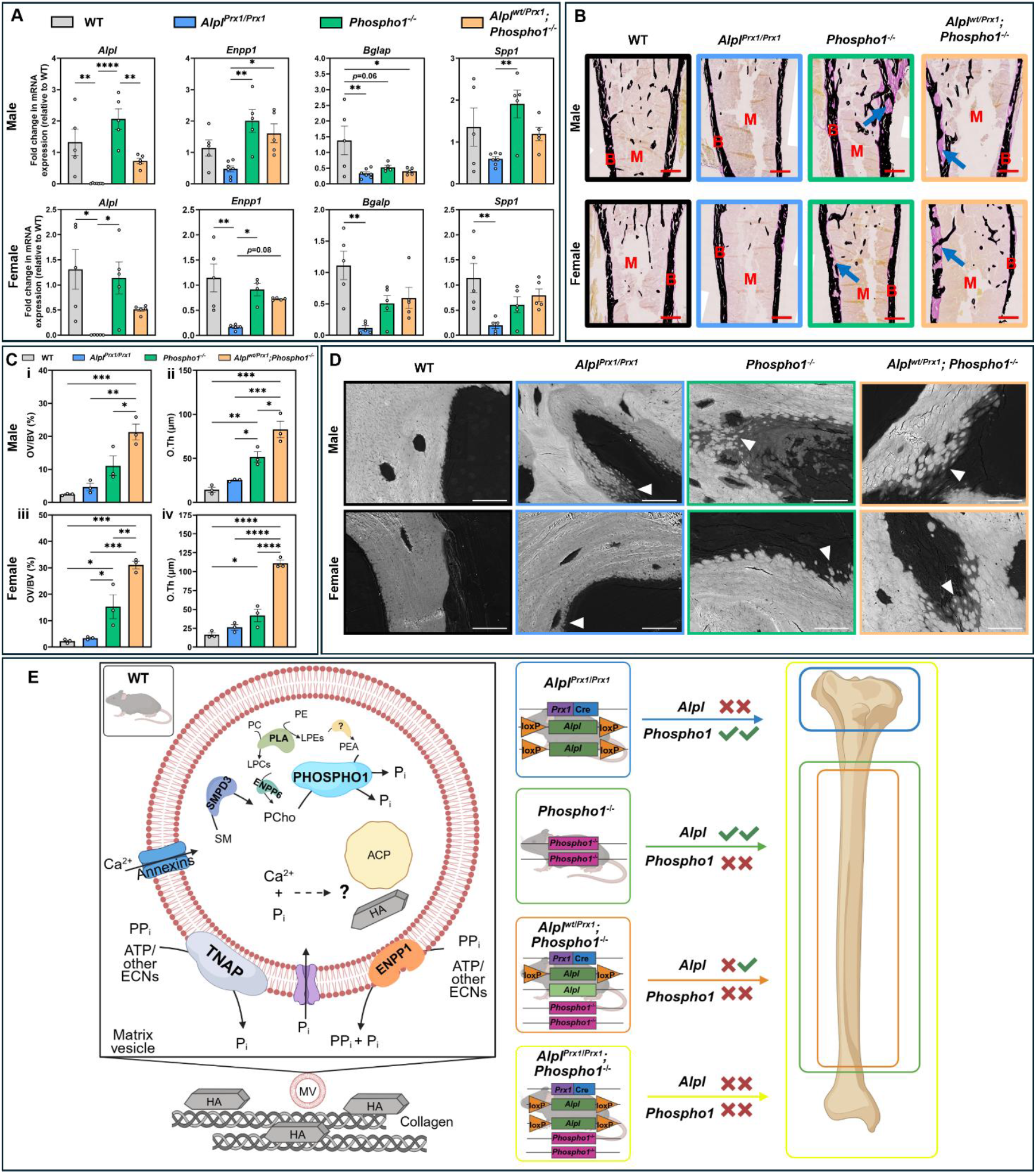
At 6-weeks of age, *Alpl*-deficient mice exhibit reduced mineralisation gene expression, yet cortical mineralisation defects are observed with *Phospho1* deletion and is compounded in *Alpl*^*wt/Prx1*^*;Phospho1*^*-/-*^ animals. (**A**) RT-qPCR of key mineralisation genes from femurs of wild-type (WT), *Alpl*^*Prx1/Prx1*^, *Phospho1*^*-/-*^ and *Alpl*^*wt/Prx1*^*;Phospho1*^*-/-*^ male and female mice (n=5-7/genotype/sex). (**B**) Representative von Kossa-stained images of the cortical region of the tibial diaphysis. Arrows indicate increased areas of osteoid in *Phospho1*^*-/-*^ and *Alpl*^*wt/Prx1*^*;Phospho1*^*-/-*^ mice. B = bone, M = bone marrow. Scale bar = 500 μm. (**C**) Quantification of (**i & iii**) osteoid volume/bone volume (OV/BV) and (**ii & iv**) osteoid thickness (O.Th) in the cortical region (n=3/genotype/sex). (**D**) Back scattered scanning electron microscopy images of the cortices. Arrowheads indicate failure of mineralisation foci to propagate within areas of hypomineralisation. Scale bar = 20 µm (**E**) Schematic showing the synergistic roles of TNAP and PHOSPHO1 in the unified model of biomineralisation and summary of our findings. Created with Biorender.com. Abbreviations: ACP: Amorphous calcium phosphate; Ca^2+^: Calcium; ECN: extracellular nucleotides; HA: hydroxyapatite; LPC: lyso-phosphatidylcholine; LPE: lyso-phosphatidylethanolamine; MV: matrix vesicle; PC: phosphatidylcholine; PCho: phosphocholine; PE: phosphatidylethanolamine; PEA: phosphoethanolamine; P_i_: inorganic phosphate; PLA: phospholipase A2; PP_i_: inorganic pyrophosphate; SM: sphingomyelin. Data are presented as mean ± SEM with points showing individual animals. *= *p* < 0.05, **= *p* < 0.01, ***= *p* < 0.001, ****= *p* < 0.0001.

## Discussion

This is the first study to overcome the perinatal lethality of global dual TNAP and PHOSPHO1 deletion in vivo to define their synergistic roles in postnatal skeletal mineralisation (6). While PHOSPHO1 and TNAP are known to liberate P_i_ for hydroxyapatite formation, our knowledge of the combined involvement of these phosphatases on postnatal skeletal structures has remained elusive. Herein, we utilised *Prx1*-*Cre* mice to target *Alpl* deletion in the early limb bud mesenchyme, including osteoblasts and chondrocytes, to generate viable *Alpl*^*Prx1/Prx1*^*;Phospho1*^*-/-*^ mice (16, 28, 29). Using a variety of multi-modal approaches, we performed full skeletal phenotyping of these mice to provide critical insights into their dual function. Our findings highlight the importance of both phosphatases in permissive biomineralisation and provide vital insights into our understanding of hypo- and hyper-mineralised pathologies, ultimately enabling future treatment strategies.

Unlike the single, stillborn, global *Alpl* and *Phospho1* double knockout mouse reported by us previously (6), our conditional knockout model survived postnatally. Here, constricting *Alpl* loss to the appendicular skeleton permitted mineralisation of the central axis. *Alpl*^*Prx1/Prx1*^*;Phospho1*^*-/-*^ mice, however, were noticeably smaller than their littermates with further skeletal phenotyping identifying a lack of mineralisation in their limbs, resembling a non-mineralised cartilaginous matrix. This confirms that permissive biomineralisation relies on the synergistic expression of these two phosphatases and adds to our mechanistic framework for understanding this process (Figure 7E) (3). Histological analysis showed distention of the growth plate cartilage into the metaphyseal region, together with an accumulation of disorganised, SOX9-positive chondrocytes. Persistent SOX9 expression suggests a suppression of hypertrophic chondrocyte-to-osteoblast transdifferentiation, which subsequently impairs metaphyseal bone structure (30). Coupled with reduced formation of the SOC and an accumulation of non-mineralised structures in the diaphyseal region, this resulted in severe limb deformation which ultimately prevented these animals from thriving post-weaning.

To align with humane practices in animal research in the UK, *Alpl*^*Prx1/Prx1*^*;Phospho1*^*-/-*^ mice were not aged above 3-weeks. To overcome this and understand the role of these phosphatases in the maturing skeleton, we additionally utilised heterozygous *Alpl*^*wt/Prx1*^*;Phospho1*^*-/-*^ animals. Even in the absence of PHOSPHO1, just a single allele of *Alpl* was sufficient to support biomineralisation, highlighting the potent effect of TNAP in this process. Surprisingly, we saw no significant differences in serum levels of PP_i_ or ALP in our 6-week-old animals, likely due to age, which is at odds with previous studies in which decreased circulating levels of ALP and increased circulating PP_i_ have been observed in mice with a *Prx*-*Cre* specific deletion of ALP (31, 32).

Our recent work in human primary osteoblasts showed that high levels of extracellular P_i_ (eP_i_) decreased expression of the phosphate transporters *Slc20a1* and *Slc20a2* (33). Similarly, eP_i_ decreased *Phospho1* and *Alpl* gene expression, whilst elevating *Enpp1* and *Ank*, implying that osteoblasts have a feedback mechanism required for them to sense and respond to altered eP_i_ levels for physiological mineralisation (33). This is broadly consistent with our findings herein in which gene expression of both *Slc20a1* and *Slc20a2* was increased with *Phospho1* deletion. Conditional ablation of *Slc20a1* in chondrocytes, in combination with global *Phospho1* deletion, worsened the skeletal phenotype than the individual gene deletions, which was marked by increased osteoid, growth plate disorganisation and reduced mechanical properties (9). However, the mechanism for P_i_ transport into the MV remains unclear (2).

Interestingly, in the global double knockout model, we previously observed some mineralisation in the axial skeleton of the single stillborn pup born which was attributed to the actions of ENPP1 conferring a compensatory role in the absence of TNAP (6). Indeed, dual deletion of *Alpl* and *Enpp1* corrects mineralisation defects observed upon single gene deletion and is thought to be due to a normalisation of PP_i_ concentrations (34). More recently, the synergistic function of TNAP and ENPP1 in ATP hydrolysis for a permissive P_i_/PP_i_ ratio has been defined (1). We found a reduction in *Enpp1* gene expression in our *Alpl*^*Prx1/Prx1*^ mice, which was normalised upon additional *Phospho1* deletion. *Smpd3* and *Chkb* expression were elevated in *Phospho1-*deficient animals, both of which are involved in the generation of the PHOSPHO1 substrate PCho and their deficiency results in a hypomineralised bone matrix (3, 35, 36).

Deletion of each phosphatase leads to altered bone development in different developmental regions of the tibia. Specifically, epiphyseal and metaphyseal changes are more often observed in *Alpl*^*Prx1/Prx1*^ mice and diaphyseal changes in *Phospho1*^*-/-*^ animals, which is further exacerbated in *Alpl*^*wt/Prx1*^*;Phospho1*^*-/-*^ heterozygous mice. Studies have reported TNAP expression was broadly expressed along the primary and secondary trabeculae, whereas PHOSPHO1 was restricted to the growth plate chondro-osseous junction, near sites of early mineral deposition, offering some potential explanation as to the spatial differences observed here (37, 38).

Our analyses in *Alpl*^*Prx1/Prx1*^ mice provide us with a model of later-onset hypophosphatasia (29). Notably, mouse strain may influence the observed phenotype, with the 129 genetic background reported to exhibit greater penetrance than the C57BL/6 one used here (29, 39). However, consistent with previous studies, we found *Alpl*^*Prx1/Prx1*^ had a failure to form SOCs with instead, large nests of hypertrophic chondrocytes (7, 40). This failure has previously been shown to be rescued upon administration of a soluble chimeric alkaline phosphatase (40). Progressive growth plate narrowing coincides with accrual of perpendicular calcified struts, herein termed growth plate bridges (41). Our 3D analysis of these bridges revealed a significant increase in bridge number in *Alpl*^*Prx1/Prx1*^ mice. We postulate this may be in response to the SOC defect, acting as a compensatory mechanism to stabilise the epiphysis in these animals. Indeed, our previous digital volume correlation and finite element modelling data support a mechanical role for growth plate bridges by revealing that their clustering is linked to elevated local stress/strain levels, as well as increased stress dissipation distal to the growth plate (41, 42). This may also have implications on the entire tibia as mechanical testing revealed that *Alpl*^*Prx1/Prx1*^ mice exhibited reduced bone mechanical properties.

Interestingly, Raman spectroscopic analysis of the epiphysis revealed sex-specific alterations in mineral and ECM composition of all genotypes, consistent with previously reported divergences in skeletal ECM organization (43). Reduced hydroxyapatite crystallinity and increased carbonate content of primary teeth from HPP patients has been reported, which opposes results observed from our *Alpl*^*Prx1/Prx1*^ mice (44). However, sex-specific differences and analysis on *Alpl*-deficient bone are yet to be investigated. Whilst our findings in *Phospho1*^*-/-*^ females appear consistent with our previous analyses (25, 43), the increased matrix carbonation in male animals is at odds. Overall, we found that dual deletion of *Alpl* and *Phospho1* resulted in changes in collagen configuration which favoured matrix carbonation and as a result compromised hydroxyapatite mineralisation, thereby reducing the mechanical integrity of the matrices (45).

In contrast to the epiphyseal and metaphyseal alterations observed with *Alpl* deletion, *Phospho1*- deficient animals had profound diaphyseal changes in cortical composition, porosity and geometry, with severe bowing observed in the distal tibia (25). Such changes were made more pronounced with the additional deletion of a single *Alpl* allele and are consistent with our previous study in 25-day-old animals in which a single *Alpl* deletion in the global knockout aggravated the skeletal phenotype of *Phospho1*^−*/*−^ mice (6). Alongside tibial shape changes, increased cortical porosity was also observed in *Phospho1*-deficient animals. Cortical porosity is a major risk factor for fragility fractures, with previous research highlighting distinct spatial variation in cortical porosity and canal density, regulated by bone-derived VEGF (46). This is associated with regional vulnerability, as is observed in the tibiofibular junction of *Phospho1*^*-/-*^ and *Alpl*^*wt/Prx1*^*;Phospho1*^*-/-*^ mice here. Such vulnerabilities are likely exacerbated by the significant mineralisation defects observed in *Phospho1*^*-/-*^ and *Alpl*^*wt/Prx1*^*;Phospho1*^*-/-*^ mice with BSSEM, akin to osteomalacia (47), and mechanical testing indeed confirmed this compromised bone phenotype in females, though remarkably no differences were observed in males.

The cortical shape changes seen in *Phospho1*^-/-^ and *Alpl*^*wt/Prx1*^*;Phospho1*^*-/-*^ mice may be linked to enlarged populations of chondrocytes in the peripheral regions of the growth plate, particularly in the medial condyle. Others have reported similar observations affecting the growth plate. Conditional deletion of ciliary Intraflagellar Transport Protein 88 (IFT88) in this region resulted in such populations, highlighting a role for the cilia in ensuring coordinated growth plate dynamics (48). Increased physiological loading inhibited peripheral growth plate ossification in WT mice, also demonstrating the sensitivity of growth plate dynamics in response to mechanical loads (48). Indeed, it is plausible that these differences in the medial peripheral growth plate may result in defective load transfer from epiphyseal regions, ultimately contributing to the differences in tibial shape of *Phospho1*^-/-^ and *Alpl*^*wt/Prx1*^*;Phospho1*^*-/-*^ mice. It is known that loading exerts significant lasting modifications in tibial shape (49) and this may be further compounded on a hypomineralised skeleton that is unable to withstand physiological load, considered a key factor in the bowing observed in *Phospho1*^-/-^ mice (25). Future work could examine the role of mechanical loading in these animals to determine the effects of TNAP and PHOSPHO1 deletion on bone mechanical properties.

In summary, our findings confirm the essential role of TNAP and PHOSPHO1 for effective biomineralisation and highlight the spatial influence they have over long bone development. By identifying synergistic phosphatase roles, this research informs enzyme replacement and gene therapy strategies, with potential to reduce healthcare costs and improve quality of life globally.

## Materials and Methods

Comprehensive methods are found in the SI Appendix.

### Animals

All tissue isolation and experimental procedures were performed in accordance with the UK Animals (Scientific Procedures) Act of 1986 and regulations set by the UK Home Office and local institutional guidelines.

Tissue-specific *Alpl*^*Prx1/Prx1*^ mice were generated using the *Prx1-Cre* transgene as previously reported (16). Similarly, the generation and characterisation of Phospho1-R74X mutant mice (Phospho1^m1Jlm^, here referred to as *Phospho1*^−*/*−^) were also previously described (6). To generate mice lacking global PHOSPHO1 and *Prx1*-conditional TNAP, *Phospho1*^−*/*−^ and *Alpl*^*Prx1/Prx1*^ were crossed to generate double mutant (*Alpl*^*Prx1/Prx1*^;*Phospho1*^−*/*−^) and *Alpl* heterozygous (*Alpl*^*wt/Prx1*^;*Phospho1*^−*/*−^) mice. Genotyping was conducted by Transnetyx using their automated genotyping service. Tissues were collected from both male and female mice at postnatal day 1 (PN1), 3- and 6-weeks of age, following euthanasia by Schedule 1. Analyses were conducted blindly to minimise the effects of subjective bias.

### Whole mount staining

PN1 and 3-week-old animals were prepared for alcian blue (0.03% w/v, 80% ethanol, 20% glacial acetic acid) and alizarin red (0.005% in 1% KOH) staining, as previously described (50).

### µCT analysis

Whole-body µCT scans were performed on PN1 and 3-week-old mice (n=3/genotype), and left tibiae from 6-week-old mice (n=5–7/genotype/sex) using a Skyscan 1172 system (Bruker) with voxel sizes of 10 µm (PN1, 40 kV), 13.5 µm (3-week, 50 kV), and 5 µm (tibiae, 40 kV). Scans were reconstructed with NRecon, aligned in DataViewer, and analysed in CTAn for tibial trabecular, epiphyseal and cortical bone analysis, as previously described (51). BMD and TMD were calibrated using standard phantoms. 3D renderings were generated in Avizo (ThermoFisher Scientific)

Growth plate bridge analysis was conducted, as previously described (41). 3D growth plate volume and thickness were manually segmented and analysed using Avizo.

Cortical porosity of the tibiofibular junction was analysed in FIJI/ImageJ and quantified using BoneJ’s Particle Analyser as previously described (46, 52).

### Three-point bending

Three-point bending was carried out on left femurs from 6-week-old animals (n=5-7/genotype/sex) using a LRX5 materials testing machine (Lloyd Instruments) fitted with a 100N load, as previously described (25). Individual load extension curves were generated to identify maximum load. Stiffness and yield were calculated from the slope of the linear region of the load extension curve using polynomial fit.

### Histological analysis

Right hindlimbs were fixed for 24 hours in 4% paraformaldehyde before being decalcified in 10% ethylenediaminetetraacetic acid for 21 days at 4°C, and wax-embedded. Coronal sections (6 μm) were stained with Haematoxylin and Eosin, Safranin O/Fast green or Toluidine blue using standard procedures. Immunohistochemical analysis was performed using anti-MMP13 and SOX9 (Table S2). The height of the growth plate proliferating and hypertrophic zones were measured at 5 different points based on established cell morphology (53).

Tartrate resistant acid phosphatase (TRAP) and von Kossa staining were conducted on methyl methacrylate (MMA) sections as described previously and sections analysed using the bone histomorphometry software, HistoMorph (54).

### Raman Spectroscopy

A total of 75 Raman spectra, presented as class means, were obtained from randomly selected locations from the tibial epiphysis in MMA-embedded sections (n=3/genotype/sex) as previously described (43, 55-57).

### BSEM

Right humeri were fixed in 4% paraformaldehyde, 2.5% glutaraldehyde (in 0.1 M sodium cacodylate) and stored in fixative at 4°C before processing and examination under back-scattered SEM (58).

### Western blotting

Femoral bone tissues were homogenised in radioimmunoprecipitation assay (RIPA) buffer containing protease inhibitors (Roche). Protein concentrations were measured by a BCA protein assay kit, separated using a 4-12% Bis-Tris protein gel, and transferred to a Polyvinylidene fluoride (PVDF) membrane. Membranes were blocked, incubated in primary antibodies overnight at 4°C, and incubated in secondary antibodies for 1 hour at room temperature (Table S2). β-actin conjugated with Horseradish Peroxidase (HRP) was incubated for 20 minutes at room temperature. The specific protein signals were detected by enhanced chemiluminescence (ECL) substrates and imaged.

### Serum PP_i_ and ALP detection

Plasma was collected by centrifugation at 2500 x*g* for 10 mins at 4°C following blood clotting for 30 mins at room temperature. PP_i_ levels were measured using a colorimetric PP_i_ assay kit (Abcam) according to the manufacturer’s instructions. ALP was detected using a colorimetric ELISA kit (Novus Biologicals), as per the manufacturer’s instructions. Absorbance was measured using a spectrophotometer (Biochrom EZ Read 2000) at 570 nm (PP_i_) or 450 nm (ALP).

### Quantitative polymerase chain reaction (qPCR) and PCR array

Left femora were cleaned of soft tissue, bone marrow removed by centrifugation and frozen at - 80°C. Bones were thawed and homogenised using a Rotor-Stator Homogenizer (IKA Ultra-Turrax T10) in ice-cold Qiazol reagent. RNA was isolated and purified using a Qiagen RNeasy mini kit, according to the manufacturer’s instructions.

RNA (2 µg) was reverse transcribed using a Qiagen QuantiTect RT kit, according to the manufacturer’s instructions. The PCR was performed using cDNA (50ng), a Brilliant II Sybr green mastermix (Agilent), and run on an Agilent AriaMx PCR machine. Gene expression data for the qPCR were normalised to *β-actin* and *Atbp5* using the geometric mean and analysed using the ΔΔCt method (59). Primer sequences can be found in Table S3.

### Statistical analysis

Analyses were performed using GraphPad Prism software (version 10.1.2). Normal distribution of data was assessed using the Shapiro-Wilk normality test. For comparison between genotypes, a one-way ANOVA followed by Tukey’s post hoc analysis was used to test for significance. Results are presented as the mean ± standard error of the mean (SEM).

For the cortical bone analysis, R software (version 4.2.2) was used to generate line graphs and perform statistical analyses. Normal distribution of data was assessed using the Shapiro-Wilk normality test. For comparison between genotypes, a one-way ANOVA followed by Tukey’s post hoc analysis was used to test for significance. Lines represent mean, with shading representing ± SEM. Colour heatmaps display the statistical significance at matched locations along the graph showing the genotype interaction and comparisons between genotypes. Red = *p*<0.001, green = *p*<0.01, yellow = *p*<0.05, grey = *p*>0.05 (not significant).

## Acknowledgments

The authors would like to thank the Bioresources Unit at the University of Brighton for the assistance with the care of animal models in this study. We also thank Prof. Denis Headon (Roslin Institute) for his discussions on the breeding strategy. We are also thankful to Dr Mark Hopkinson (Royal Veterinary College) and the Histology SRF at the University of Liverpool for their technical assistance. The authors would like to acknowledge the UKRI Medical Research Council (MRC) for funding to KAS (MR/V033506/1 & MR/R022240/2), the Biotechnology and Biological Sciences Research Council (BBSRC) for Institute Strategic Programme Grant Funding to CF/SNJ (BBS/E/D/10002071 & BBS/E/RL/230001C) and for supporting LAS via a Discovery Fellowship (BB/X009904/1), and NIDCR (R01DE032334) for funding BLF. For the purpose of open access, the authors have applied a Creative Commons Attribution (CC-BY) license to any Author Accepted Manuscript version arising from this submission.

