## Supplementary info for "TNAP and PHOSPHO1 function synergistically to afford critical control over the mineralisation of the postnatal murine skeleton"

**Supporting information Appendix**

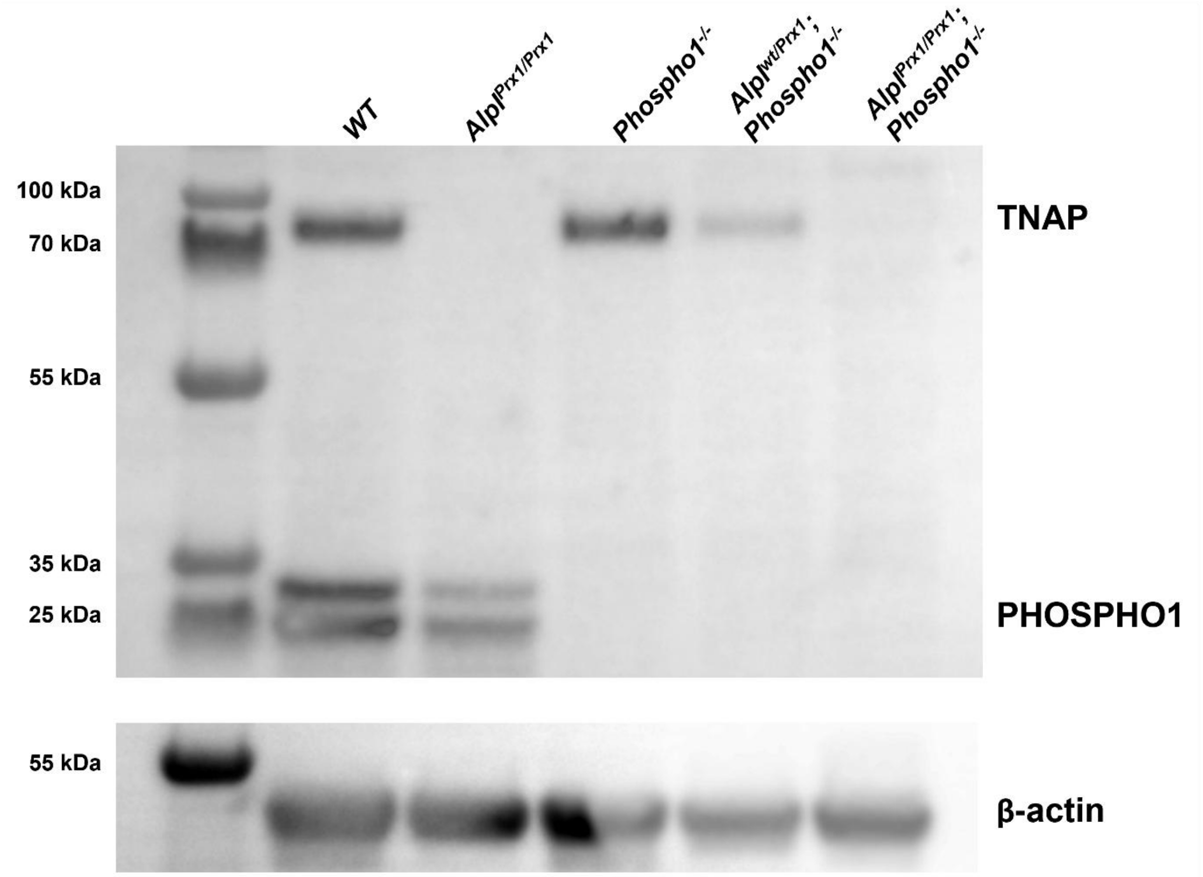


**Supplementary Figure 1. Confirmation of knockout in femoral samples.** Western blot on protein lysates of wild-type (WT), $Alpl^{Prx1/Prx1}$, $Phospho1^{-/-}$, $Alpl^{wt/Prx1}$;$Phospho1^{-/-}$, and $Alpl^{Prx1/Prx1}$;$Phospho1^{-/-}$ mice shows the loss of PHOSPHO1 expression in $Phospho1^{-/-}$ containing genotypes and loss of TNAP expression in $Alpl^{Prx1/Prx1}$ containing genotypes, normalised to β-actin. $Alpl^{wt/Prx1}$;$Phospho1^{-/-}$ shows a reduction in TNAP expression compared to WT and $Phospho1^{-/-}$.

**Supporting information Appendix**

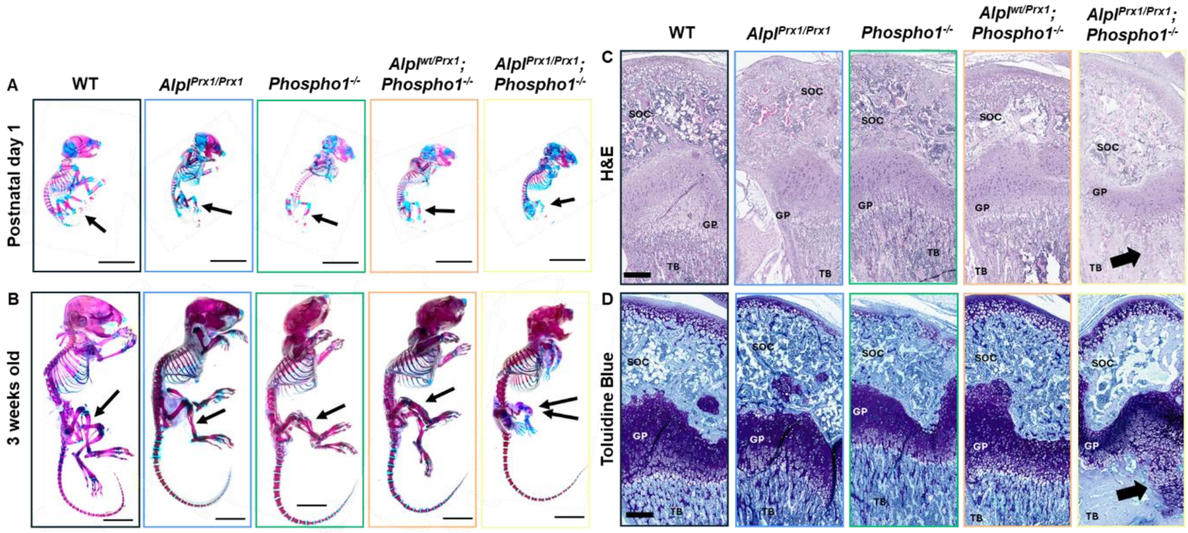


**Supplementary Figure 2. Whole mount imaging of entire skeletons at PN1 and 3-weeks-old, and histological analysis of 3-week-old mice.** Representative images of wild-type (WT), *Alpl*$^{Prx1/Prx1}$, *Phospho1*$^{-/-}$, *Alpl*$^{wt/Prx1}$;*Phospho1*$^{-/-}$, and *Alpl*$^{Prx1/Prx1}$;*Phospho1*$^{-/-}$ mice at (**A**) postnatal day 1 and (**B**) 3 weeks old. Arrows indicate mineralisation in all genotypes except for *Alpl*$^{Prx1/Prx1}$;*Phospho1*$^{-/-}$ double knockout mice. Scale bar = 1cm. (**C**) Representative images of Haematoxylin/Eosin and (**D**) Toluidine Blue staining of femurs from each genotype at 3 weeks of age. Arrows indicate enlarged growth plate extending into the metaphyseal region. Scale bar = 200 µm. GP = growth plate, SOC = secondary ossification centres, TB = metaphyseal trabecular bone.

**Supporting information Appendix**

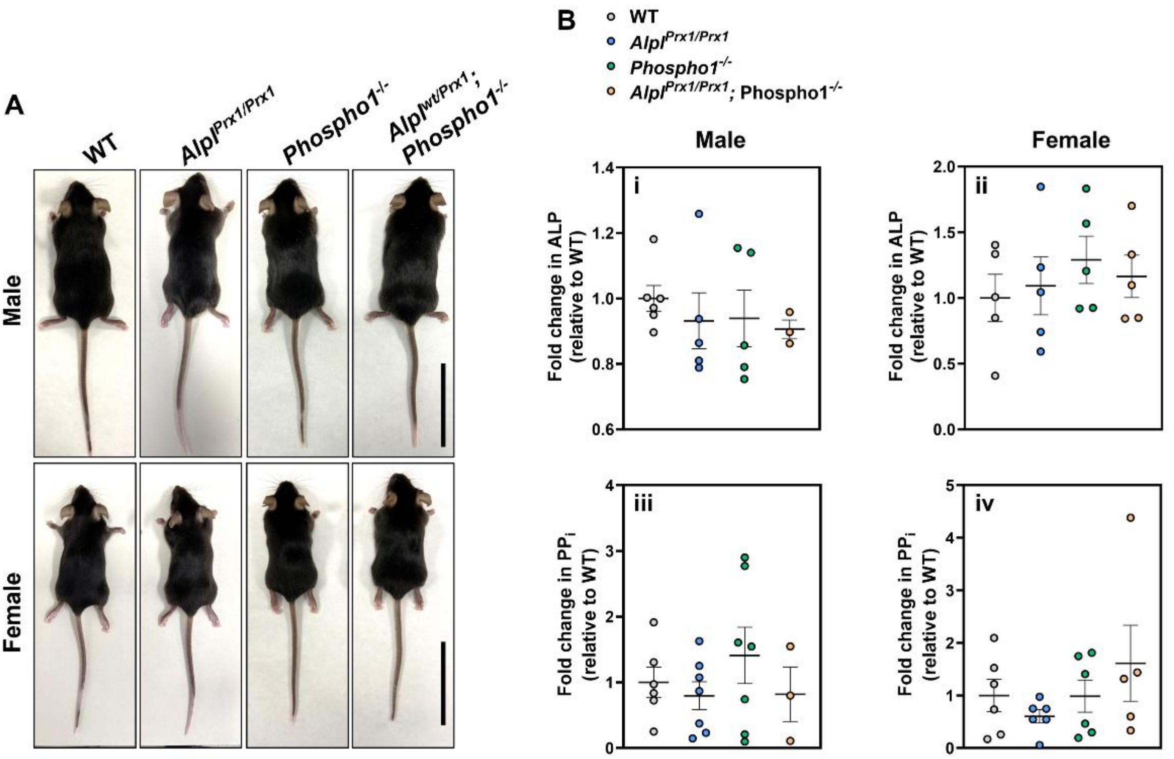


**Supplementary Figure 3. No difference in gross morphology or serum biochemistry of 6-week-old animals.** (**A**) Representative images of wild-type (WT), *Alpl*$^{Prx1/Prx1}$, *Phospho1*$^{-/-}$, *Alpl*$^{wt/Prx1}$;*Phospho1*$^{-/-}$, and *Alpl*$^{Prx1/Prx1}$;*Phospho1*$^{-/-}$ mice. Scale bar = 2cm. (**B**) Serum biochemistry analysis for **(i & ii**) ELISA detecting alkaline phosphatase (ALP) and (**iii & iv**) colorimetric assay detecting pyrophosphate ($PP_i$) in males and females, respectively. Data are presented as mean ± SEM with points showing individual animals (n=3-7/genotype/sex).

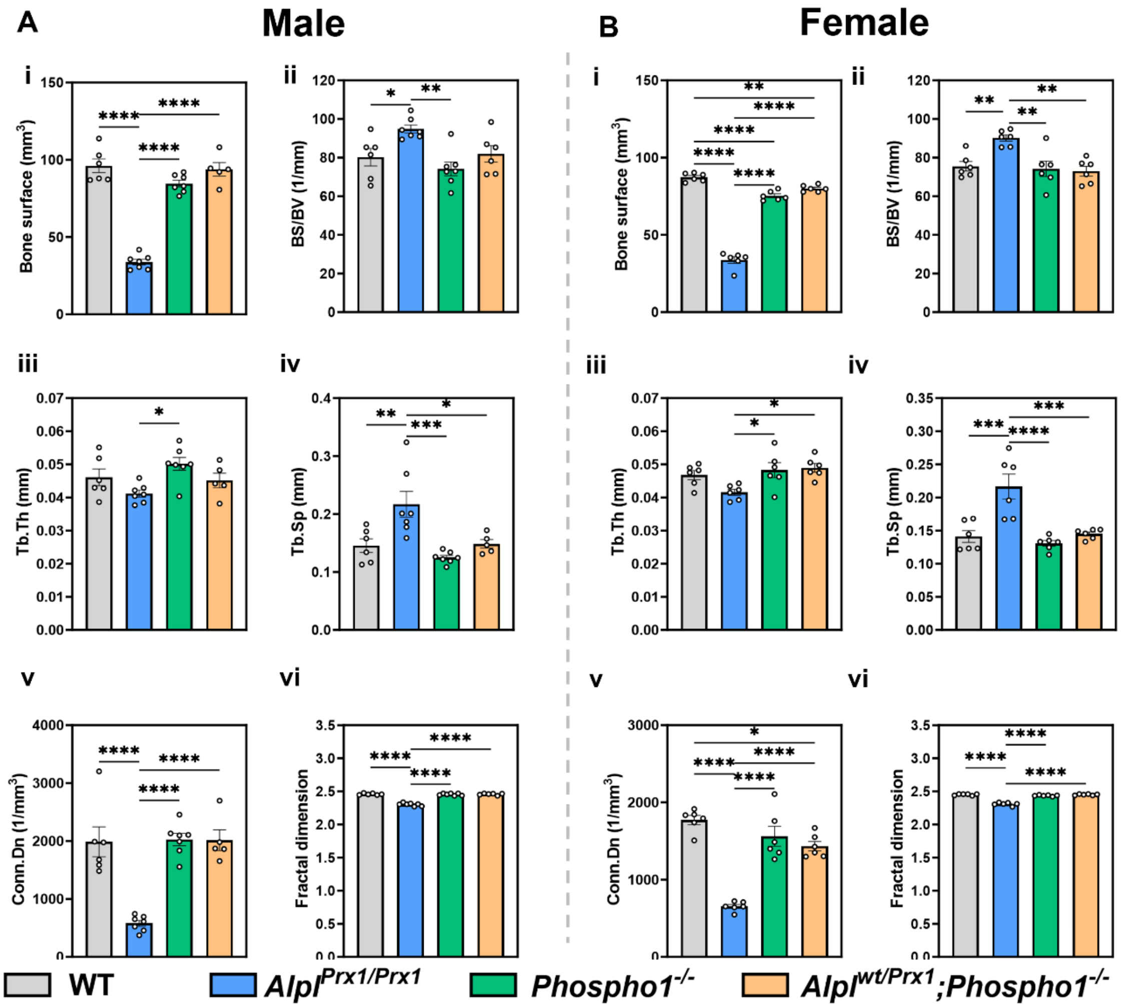


**Supplementary Figure 4. Additional trabecular parameters from the epiphyseal region of 6-week-old male and female mice.** µCT analysis of trabecular parameters from wild-type (WT), *Alpl*$^{Prx1/Prx1}$, *Phospho1*$^{-/-}$, *Alpl*$^{wt/Prx1}$;*Phospho1*$^{-/-}$, and *Alpl*$^{Prx1/Prx1}$;*Phospho1*$^{-/-}$ mice in (**A**) males and (**B**) females, including (**i**) bone surface, (**ii**) bone surface to bone volume fraction (BS/BV), (**iii**) trabecular thickness (Tb.Th), (**iv**) trabecular spacing (Tb.Sp), (**v**) connectivity density (Conn. Dn), and (**vi**) fractal dimension. Data are presented as mean ± SEM with points showing individual animals (n=5-7/genotype/sex). *= $p < 0.05$, **= $p < 0.01$, ***= $p < 0.001$, ****= $p < 0.0001$.

**Supporting information Appendix**

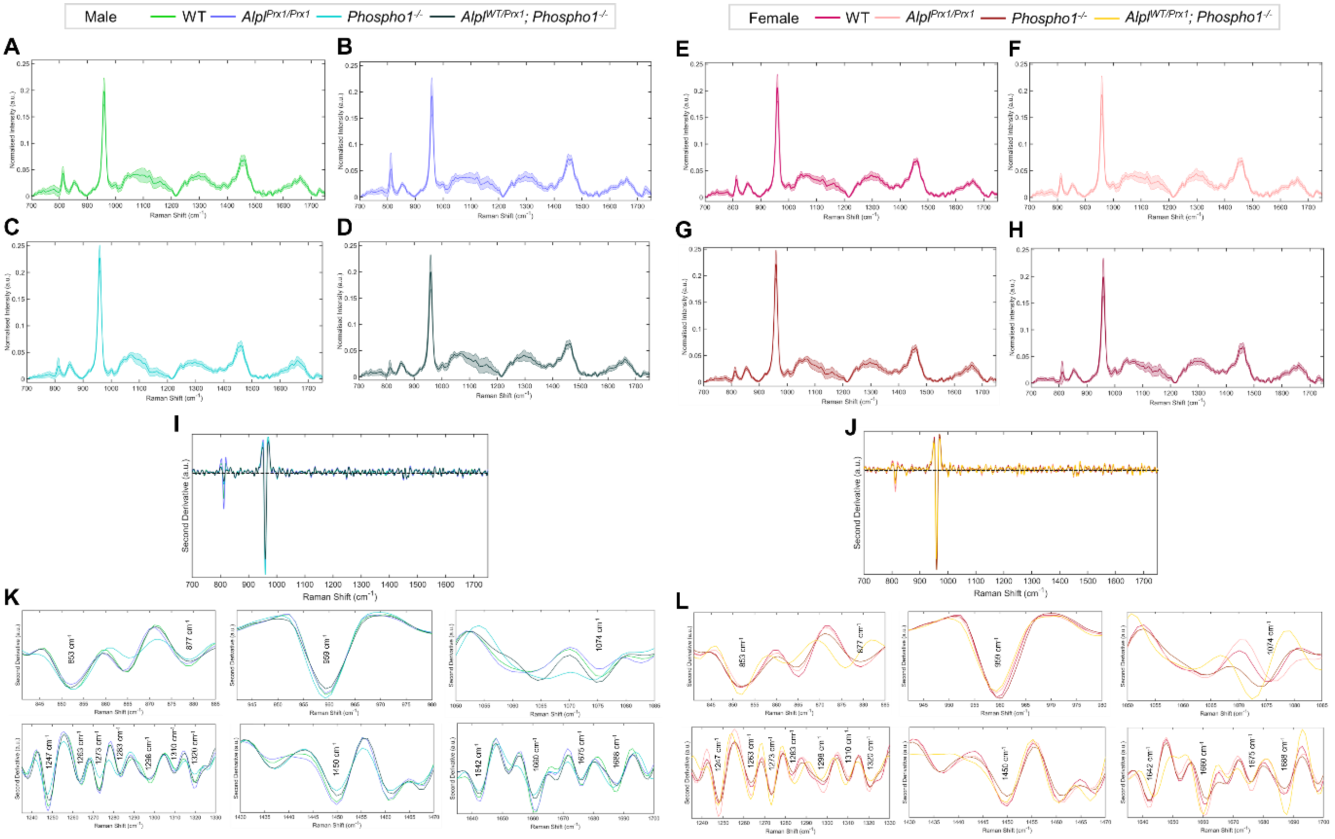


**Supplementary Figure 5. Raman spectra for 6-week-old mice.** Raman spectra collected within the fingerprint region from 700 cm-1 to 1750 cm-1 were obtained from the tibial epiphyses of (**A&E**) wild-type (WT), (**B&F**) *Alpl*$^{Prx1/Prx1}$, (**C&G**) *Phospho1*$^{-/-}$, and (**D&H**) *Alp*$^{wt/Prx1}$;*Phospho1*$^{-/-}$ male and female mice, respectively (n=3/sex/genotype). Presented as vector normalised class means spectra (n=75 individual spectra) ±SD. Second derivative calculation (male, **I** and female, **J**) facilitated the identification of Raman bands corresponding to extracellular matrix and mineral components, shown in zoomed second derivative spectra (male, **K** and female, **L**) which were analysed by spectral deconvolution.

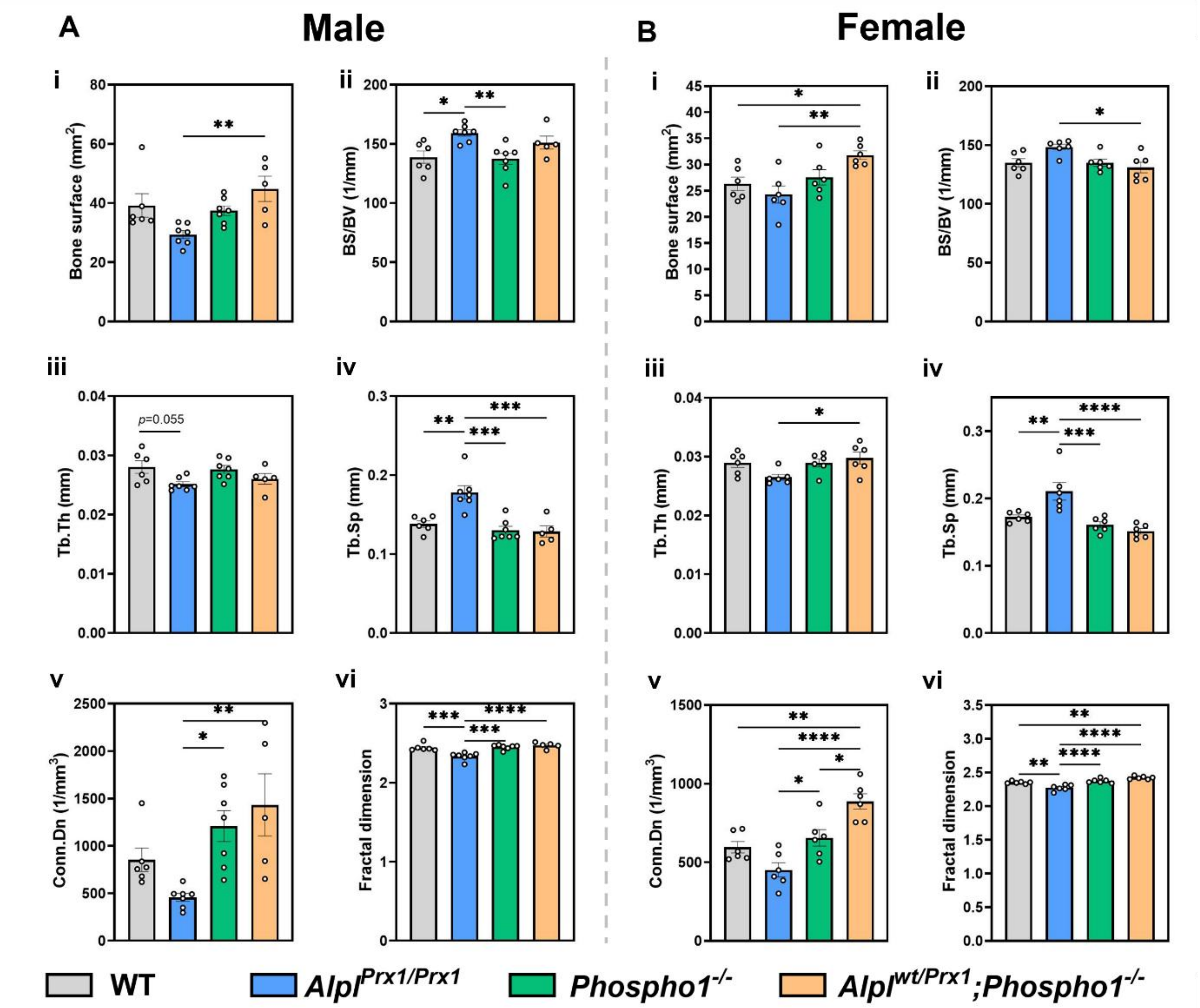


**Supplementary Figure 6. Additional trabecular parameters from the metaphyseal region of 6-week-old mice.** µCT analysis of trabecular parameters from wild-type (WT), *Alpl*$^{Prx1/Prx1}$, *Phospho1*$^{-/-}$, *Alpl*$^{wt/Prx1}$;*Phospho1*$^{-/-}$, and *Alpl*$^{Prx1/Prx1}$;*Phospho1*$^{-/-}$ mice in (**A**) males and (**B**) females, including (**i**) bone surface, (**ii**) bone surface to bone volume fraction (BS/BV), (**iii**) trabecular thickness (Tb.Th), (**iv**) trabecular spacing (Tb.Sp), (**v**) connectivity density (Conn. Dn), and (**vi**) fractal dimension. Data are presented as mean ± SEM with points showing individual animals (n=5-7/genotype/sex). *= $p < 0.05$, **= $p < 0.01$, ***= $p < 0.001$, ****= $p < 0.0001$.

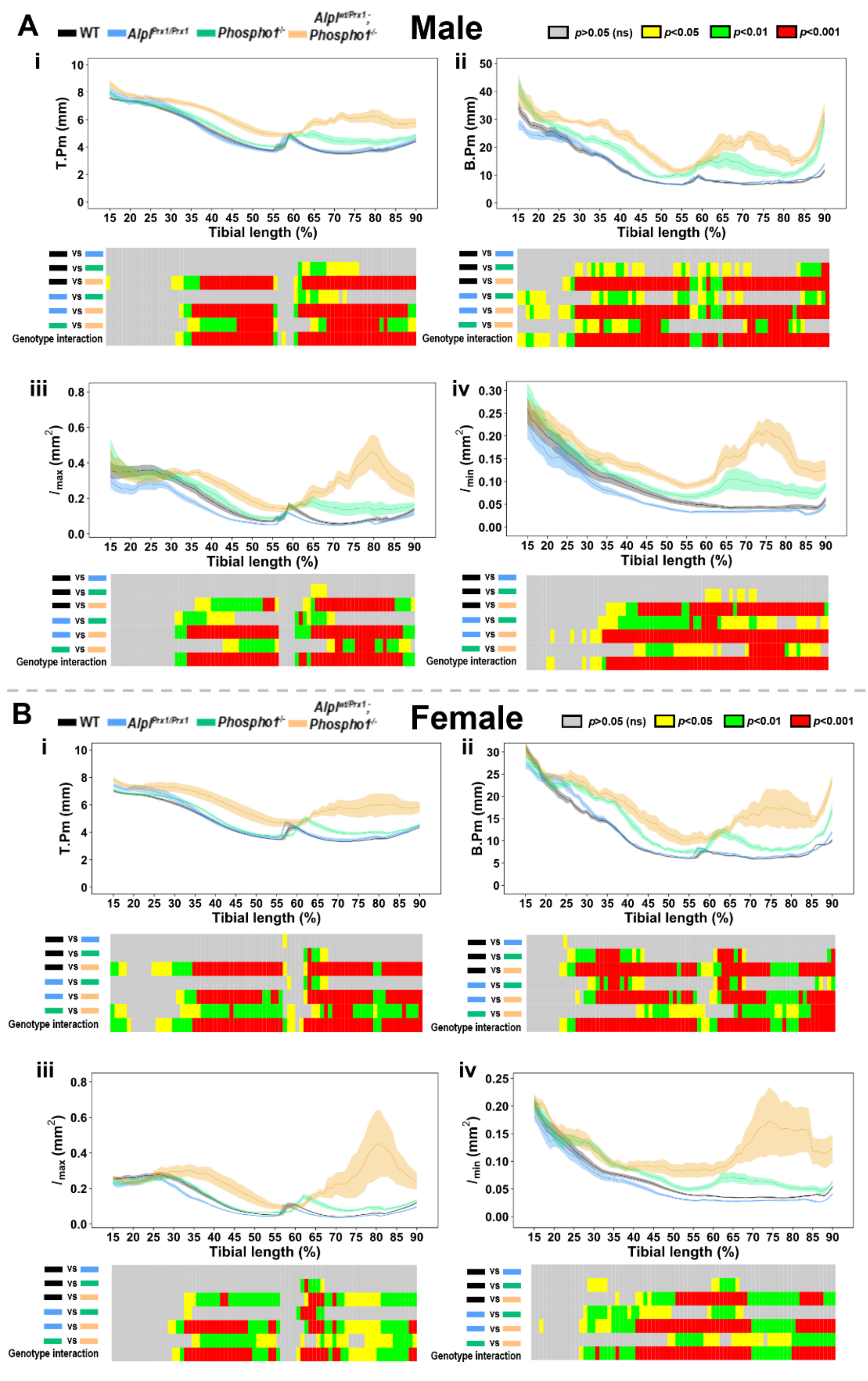


**Supplementary Figure 7. Additional cortical parameters in 6-week-old mice.** Measurement and statistical analysis heat map in (**A**) male and (**B**) female mice for (**i**) tissue perimeter (T.Pm), (**ii**) bone perimeter (B.Pm), (**iii**) maximum second moments of inertia ($I_{max}$) and, (**iv**) minimum second moments of inertia ($I_{min}$). Line graphs represent mean ± SEM for wild-type (WT; black), *Alpl*$^{Prx1/Prx1}$ (blue), *Phospho1*$^{-/-}$ (green) and *Alpl*$^{wt/Prx1}$;*Phospho1*$^{-/-}$ (orange) mice (n=5-7/genotype/sex). Graphical heat map summarises statistical differences at specific matched locations along the tibial length (15–90%). Red = $p < 0.001$, green = $p < 0.01$, yellow = $p < 0.05$, grey = $p > 0.05$ (not significant).

**Supporting information Appendix**

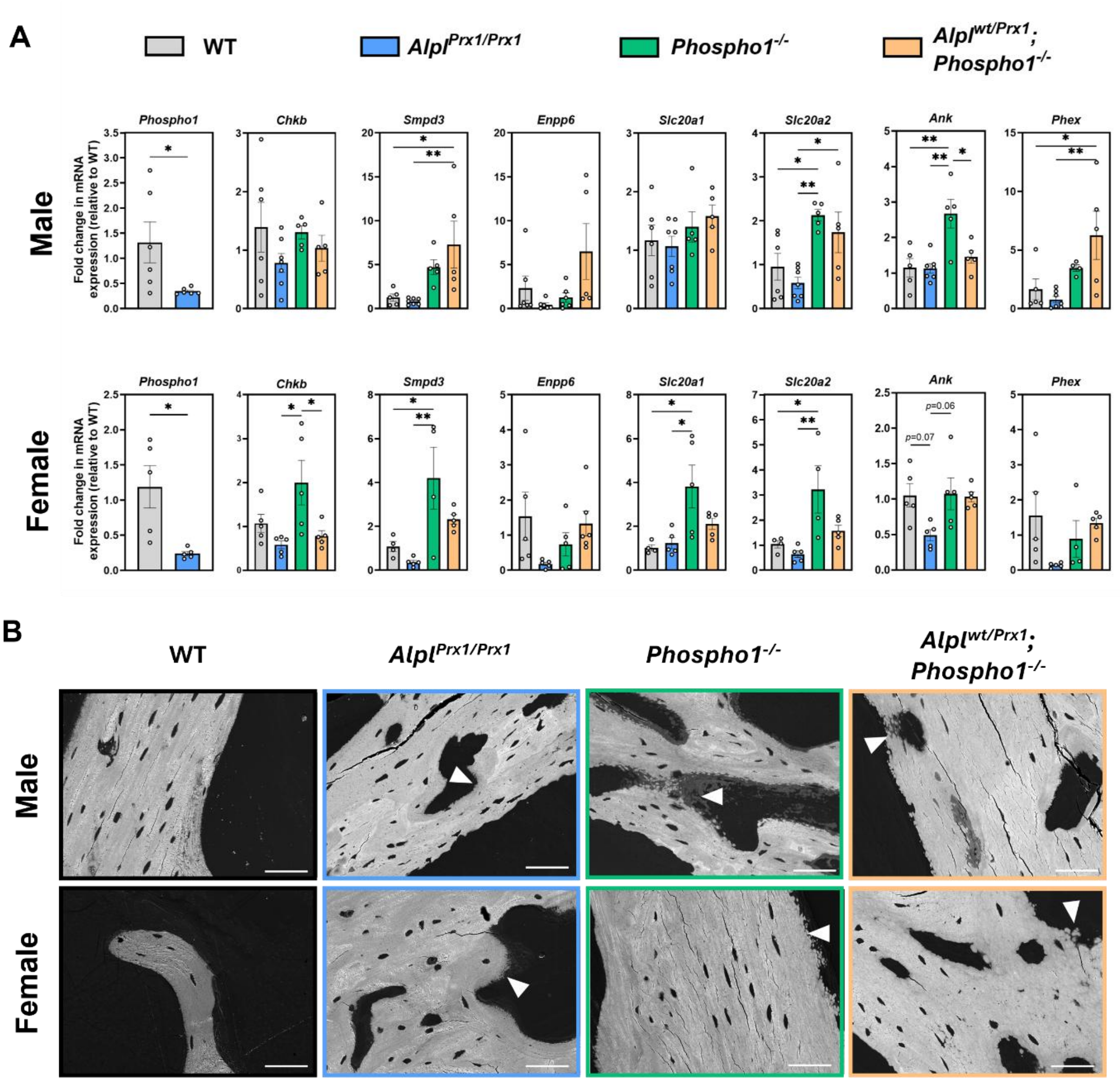


**Supplementary Figure 8. Additional genes analysed by RT-qPCR & BSEM images.** (**A**) RT-qPCR of additional genes from the femurs of wild-type (WT), *Alpl*$^{Prx1/Prx1}$, *Phospho1*$^{-/-}$, *Alpl*$^{wt/Prx1}$;*Phospho1*$^{-/-}$, and *Alpl*$^{Prx1/Prx1}$;*Phospho1*$^{-/-}$ mice, in males and females. Data are presented as mean ± SEM with points showing individual animals (n=5-7/genotype/sex). *= $p < 0.05$, **= $p < 0.01$. (**B**) Back scattered scanning electron microscopy images of the cortices. Arrowheads indicate failure of mineralisation foci to propagate within areas of hypomineralisation. Scale bar = 50 μm.

**Supplementary Table 1**: Gross morphological measurements for 6-week-old male and female mice. Data represented as mean ± SEM for $n$=5-7/genotype/sex. * = $p < 0.05$, ** = $p < 0.01$, *** = $p < 0.001$, **** = $p < 0.0001$.

| **Male** | | | | |
|---|---|---|---|---|
| | **WT** | $Alpl^{Prx1/Prx1}$ | $Phospho1^{-/-}$ | $Alpl^{wt/Prx1}$; $Phospho1^{-/-}$ |
| **Weight (g)** | 24.39 ±0.78 | 23.15 ±0.66 | 21.11 ±0.51 | 22.07 ±0.60 |
| Multiple comparisons test results (Tukey's HSD) | | | | |
| WT vs. $Alpl^{Prx1/Prx1}$ | ns | | | |
| WT vs. $Phospho1^{-/-}$ | ** | | | |
| WT vs. $Alpl^{wt/Prx1}$;$Phospho1^{-/-}$ | ns | | | |
| $Alpl^{Prx1/Prx1}$ vs. $Phospho1^{-/-}$ | ns | | | |
| $Alpl^{Prx1/Prx1}$ vs. $Alpl^{wt/Prx1}$;$Phospho1^{-/-}$ | ns | | | |
| $Phospho1^{-/-}$ vs. $Alpl^{wt/Prx1}$;$Phospho1^{-/-}$ | ns | | | |
| | **WT** | $Alpl^{Prx1/Prx1}$ | $Phospho1^{-/-}$ | $Alpl^{wt/Prx1}$; $Phospho1^{-/-}$ |
| **Nose-to-tail length (cm)** | 16.48 ±0.16 | 16.80 ±0.16 | 15.64 ±0.13 | 15.54 ±0.244 |
| Multiple comparisons test results (Tukey's HSD) | | | | |
| WT vs. $Alpl^{Prx1/Prx1}$ | ns | | | |
| WT vs. $Phospho1^{-/-}$ | ** | | | |
| WT vs. $Alpl^{wt/Prx1}$;$Phospho1^{-/-}$ | ** | | | |
| $Alpl^{Prx1/Prx1}$ vs. $Phospho1^{-/-}$ | *** | | | |
| $Alpl^{Prx1/Prx1}$ vs. $Alpl^{wt/Prx1}$;$Phospho1^{-/-}$ | *** | | | |
| $Phospho1^{-/-}$ vs. $Alpl^{wt/Prx1}$;$Phospho1^{-/-}$ | ns | | | |
| | **WT** | $Alpl^{Prx1/Prx1}$ | $Phospho1^{-/-}$ | $Alpl^{wt/Prx1}$; $Phospho1^{-/-}$ |
| **Bone length (mm)** | 16.82 ±0.10 | 16.19 ±0.10 | 14.50 ±0.31 | 13.32 ±0.32 |
| Multiple comparisons test results (Tukey's HSD) | | | | |
| WT vs. $Alpl^{Prx1/Prx1}$ | ns | | | |
| WT vs. $Phospho1^{-/-}$ | **** | | | |
| WT vs. $Alpl^{wt/Prx1}$;$Phospho1^{-/-}$ | **** | | | |
| $Alpl^{Prx1/Prx1}$ vs. $Phospho1^{-/-}$ | **** | | | |
| $Alpl^{Prx1/Prx1}$ vs. $Alpl^{wt/Prx1}$;$Phospho1^{-/-}$ | **** | | | |
| $Phospho1^{-/-}$ vs. $Alpl^{wt/Prx1}$;$Phospho1^{-/-}$ | * | | | |
| **Female** | | | | |
| | **WT** | $Alpl^{Prx1/Prx1}$ | $Phospho1^{-/-}$ | $Alpl^{wt/Prx1}$; $Phospho1^{-/-}$ |
| **Weight (g)** | 19.51 ±0.37 | 17.86 ±0.82 | 17.22 ±0.53 | 18.13 ±0.46 |
| Multiple comparisons test results (Tukey's HSD) | | | | |
| WT vs. $Alpl^{Prx1/Prx1}$ | ns | | | |
| WT vs. $Phospho1^{-/-}$ | * | | | |
| WT vs. $Alpl^{wt/Prx1}$;$Phospho1^{-/-}$ | ns | | | |
| $Alpl^{Prx1/Prx1}$ vs. $Phospho1^{-/-}$ | ns | | | |
| $Alpl^{Prx1/Prx1}$ vs. $Alpl^{wt/Prx1}$;$Phospho1^{-/-}$ | ns | | | |
| $Phospho1^{-/-}$ vs. $Alpl^{wt/Prx1}$;$Phospho1^{-/-}$ | ns | | | |
| | **WT** | $Alpl^{Prx1/Prx1}$ | $Phospho1^{-/-}$ | $Alpl^{wt/Prx1}$; $Phospho1^{-/-}$ |
| **Nose-to-tail length (cm)** | 16.05 ±0.12 | 15.96 ±0.22 | 15.07 ±0.22 | 15.57 ±0.13 |
| Multiple comparisons test results (Tukey's HSD) | | | | |
| WT vs. $Alpl^{Prx1/Prx1}$ | ns | | | |

**Supporting information Appendix**

| | | | | |
|---|---|---|---|---|
| WT vs. $Phospho1^{-/-}$ | | ** | | |
| WT vs. $Alpl^{wt/Prx1};Phospho1^{-/-}$ | | ns | | |
| $Alpl^{Prx1/Prx1}$ vs. $Phospho1^{-/-}$ | | * | | |
| $Alpl^{Prx1/Prx1}$ vs. $Alpl^{wt/Prx1};Phospho1^{-/-}$ | | ns | | |
| $Phospho1^{-/-}$ vs. $Alpl^{wt/Prx1};Phospho1^{-/-}$ | | ns | | |
| | **WT** | $\boldsymbol{Alpl^{Prx1/Prx1}}$ | $\boldsymbol{Phospho1^{-/-}}$ | $\boldsymbol{Alpl^{wt/Prx1};}$ $\boldsymbol{Phospho1^{-/-}}$ |
| **Bone length (mm)** | 16.46 ±0.09 | 15.68 ±0.13 | 14.21 ±0.15 | 13.18 ±0.28 |
| Multiple comparisons test results (Tukey's HSD) | | | | |
| WT vs. $Alpl^{Prx1/Prx1}$ | | * | | |
| WT vs. $Phospho1^{-/-}$ | | **** | | |
| WT vs. $Alpl^{wt/Prx1};Phospho1^{-/-}$ | | **** | | |
| $Alpl^{Prx1/Prx1}$ vs. $Phospho1^{-/-}$ | | **** | | |
| $Alpl^{Prx1/Prx1}$ vs. $Alpl^{wt/Prx1};Phospho1^{-/-}$ | | **** | | |
| $Phospho1^{-/-}$ vs. $Alpl^{wt/Prx1};Phospho1^{-/-}$ | | ** | | |

**Supplementary Table 2**: Western blot and immunohistochemistry (IHC) antibodies and details

| **Antibody** | **Source** | **Concentration** | **Company & catalogue number** | **Application** |
|---|---|---|---|---|
| β-actin | Conjugated with HRP | 1:1000 | Cell Signaling Technology | Western blot |
| MMP13 | Rabbit | 1:200 | Abcam (ab39012) | IHC |
| PHOSPHO1 | Human | 1:250 | Bio-Rad (HCA093) | Western blot |
| SOX9 | Rabbit | 1:500 | Merck (AB5535) | IHC |
| TNAP | Human | 1:500 | R&D Systems | Western blot |
| anti-Human IgG (H/L): HRP | Goat | 1:1000 | Rio-Rad (5172-2504) | Western blot (secondary) |

**Supplementary Table 3.** Primer sequences used for RT-qPCR

| **Gene** | **Forward sequence** | **Reverse sequence** |
|---|---|---|
| ***Alpl*** | GGGACGAATCTCAGGGTACA | AGTAACTGGGGTCTCTCTCTTT |
| ***Ank*** | TCGCTGCCTTCCCTTTTATG | GGTGACTGTGAAGCAAAATGG |
| ***Atpb5*** | ATGCCACTTCCAAGGTAGCG | GCAACAGTCAGACCAGTCAGA |
| ***βactin*** | GATGTGGCACCACACCTTCT | GGGGTGTTGAAGGTCTCAAA |
| ***Bglap*** | CCGGGAGCAGTGTGAGCTTA | TAGATGCGTTTGTAGGCGGTC |
| ***Chkb*** | GCAAGACCCACGGACTACC | CAGGGAGTACCGACTGATCTC |
| ***Enpp1*** | GCTAATCATCAGGAGGTCAAG | CTGGTAGAATCCCGTCAATC |
| ***Enpp6*** | GTAGTCATCTTGGACCCTCTCATACTG | GTGTGAGCTCTTACATGTGGACAGA |
| ***Phex*** | CTAACCACCACTCCCACTT | CCAATAGACTCCAACCTGAAGA |
| ***Phospho1*** | GCCTGCGTGCCACCTAT | CCTGCTCACCCAGGTACTTAAA |
| ***Slc20a1*** | GGCTCAGGTGTAGTGACCCT | CACATCTATCAAGCCGTTCC |
| ***Slc20a2*** | CCATCGGCTTCTCACTCGT | AAACCAGGAGGCGACAATCT |
| ***Smpd3*** | ACACGACCCCCTTTCCTAATA | GGCGCTTCTCATAGGTGGTG |
| ***Spp1*** | GAGAGCCAGGAGAGTGCCGA | GCTTTGGAACTTGCTTGACTATCG |

**Supporting information Appendix**

### Methods

*Reagents*

Unless otherwise stated, all reagents were obtained from ThermoFisher Scientific, and chemicals were purchased from Merck.

*Animals*

All tissue isolation and experimental procedures were performed in accordance with the UK Animals (Scientific Procedures) Act of 1986 and regulations set by the UK Home Office and local institutional guidelines at the University of Brighton.

Tissue-specific $\textit{Alpl}^{\textit{Prx1/Prx1}}$ mice were generated using the *Prx1-Cre* transgene as previously reported (1). Similarly, the generation and characterisation of Phospho1-R74X mutant mice ($\text{Phospho1}^{\text{m1Jlm}}$, here referred to as $\textit{Phospho1}^{-/-}$) were also previously described (2). To generate mice lacking global PHOSPHO1 and *Prx1*-conditional TNAP, $\textit{Phospho1}^{-/-}$ and $\textit{Alpl}^{\textit{Prx1/Prx1}}$ were crossed to generate double mutant ($\textit{Alpl}^{\textit{Prx1/Prx1}}$;$\textit{Phospho1}^{-/-}$) and *Alpl*-heterozygous ($\textit{Alpl}^{\textit{wt/Prx1}}$;$\textit{Phospho1}^{-/-}$) mice. Genotyping was conducted by Transnetyx using their automated genotyping service. Tissues were collected from both male and female mice at postnatal day 1 (PN1), 3- and 6- weeks of age, following euthanasia by Schedule 1. Analyses were conducted blindly to minimise the effects of subjective bias.

*Whole mount staining*

PN1 and 3-week-old animals were skinned, eviscerated and fixed in 95% ethanol overnight, as previously described (3). Acetone (100%) was replaced for 1-2 days. Preparations were incubated in 0.03% alcian blue (w/v), 80% ethanol, 20% glacial acetic acid for 1-3 days, before destaining in two changes of 70% ethanol and incubating in 95% ethanol overnight. Samples were pre-cleared in 1% potassium hydroxide (KOH) solution overnight at 4°C and incubated in 0.005% alizarin red (in 1% KOH) at 4°C for 1-3 days. Specimens were cleared in 1% KOH and stored in 50% glycerol:50% KOH (1%) before imaging.

*MicroCT (µCT) analysis*

Whole body scans of PN1 (n=3/genotype) and 3-week-old animals (n=3/genotype), and 6-week-old left tibiae (n=5-7/genotype/sex) were performed using a Skyscan 1172 X-Ray microtomograph (Bruker). Tomographic scans (200 µA, 0.5 mm aluminium filter, 0.6° rotation angle) were acquired with a voxel size of 10 µm (40 kV) for PN1 animals; 13.5 µm (50 kV) for 3-week-old animals; 5 µm for 6-week-old tibiae (40 kV). The projection images were reconstructed using NRecon software (Bruker, version 1.7.4.6). Each dataset was orientated and aligned in DataViewer (Bruker, version 1.5.6.2). Region of interest segmentation was performed in CTAn software (Skyscan).

In PN1 and 3-week-old mice, bone surface area of the entire skeleton was performed on binarised images using a minimum threshold of 80. 3D measurements were obtained in CTAn. In tibiae, 3D analysis of the trabecular bone was performed using a volume of interest of 5% of the total bone length, as previously described (4). This region extended distally from the bottom of the growth plate where the primary spongiosa is no longer visible towards the diaphysis. Analysis of the epiphysis included the most proximal part of the tibia to the unmineralised growth plate. A binarised image with a minimum threshold set to 85 was used across all datasets and 3D measurements taken in CTAn. BMD measurements were calibrated with appropriate phantoms. Cortical bone was analysed using a 2D whole-bone approach to visualise changes across the entire tibia (4). To ensure exclusion of the trabecular bone, the proximal 15% and distal 10% of the tibial length were excluded. Slice by slice measurements were created in CTAn using binarised images with a minimum threshold set to 80. Volumetric tissue mineral density (TMD) measurements were calibrated using appropriate phantoms. 3D-reconstructed images for the entire skeleton, and the cortical and trabecular regions of the tibia were created using Avizo software (ThermoFisher Scientific).

Growth plate bridge analysis was conducted as previously described whereby the central points of all bony bridges were identified and projected on the tibial joint surface to determine spatial distribution (5). To evaluate 3D growth plate volume ($\mu m^3$) and thickness (µm, parallel plate model), proximal tibial growth plates were manually segmented using Avizo. To generate false-colour thickness heatmaps, segmentations were resampled, and the Avizo 'thickness map' module with the 'physics' Lookup Table applied.

To further examine cortical porosity, tibial µCT scans (500 slices from the tibiofibular junction) were processed in FIJI ImageJ. Thresholding produced binary images, and 'Keep Largest Region' isolated the tibia. A filled mask minus the marrow cavity yielded a cortical tissue mask. Intracortical canals were identified by subtracting the binary tibia image, excluding osteocyte lacunae due to resolution limits (6). BoneJ's 'Particle Analyser' quantified cortical porosity (7).

*Three-point bending*

Left femurs from 6-week-old animals (n=5-7/genotype/sex) were cleaned of soft tissue and frozen in distilled water at -20°C. Three-point bending was carried out on thawed femurs using a LRX5 materials testing machine (Lloyd Instruments) fitted with a 100N load. The span was fixed at 10mm and bones loaded with a crosshead speed of 1mm/min until failure (8). Load extension curves were generated to identify maximum load. Stiffness and yield were calculated from the slope of the linear region of the load extension curve using polynomial fit.

*Histological analysis*

Right hindlimbs were fixed for 24 hours in 4% paraformaldehyde before being decalcified in 10% ethylenediaminetetraacetic acid for 21 days at 4°C, and wax-embedded. Coronal sections (6 μm) were stained with Haematoxylin and Eosin, Safranin O/Fast green and Toluidine blue using standard procedures. Immunohistochemical analysis was performed using anti-MMP13 and SOX9 (Table S2). Briefly, sections were dewaxed in xylene and rehydrated. For antigen retrieval, sections were incubated at 37°C for 30 mins in 1mg/ml trypsin (MMP13) or 10mM sodium citrate at 60°C for 90 mins (SOX9). Endogenous peroxidases were blocked by treatment with 0.3% $H_2O_2$ in methanol for 30 mins at room temperature. The Vectastain ABC universal detection kit (Vector Laboratories) was used according to the manufacturer's instructions, with diaminobenzidine (DAB) solution used to detect the location of antigens. The sections were finally dehydrated and mounted in DPX. The height of the growth plate proliferating and hypertrophic zones were measured at 5 different points based on established cell morphology (9).

For methyl methacrylate (MMA) histology, samples were dehydrated through an alcohol series and two changes of xylene at 4°C using a Leica ASP 300 tissue processor. The samples were infiltrated at 4°C under vacuum with a mixture of 88.99% MMA, 10% di-butyl phthalate, 1% Perkadox 16 and 0.01% Tinogard® TT for 5 days. After infiltration, the samples were placed in Teflon embedding blocks and left to polymerise in a water bath at room temperature for a minimum of 2 days. The lids were subsequently removed and embedding rings were attached using Technovit 3040. Blocks were left to harden for at least 2 days before sectioning at 5μm using a Leica RM2265 motorised microtome fitted with a tungsten steel D-profile knife. The sections were mounted on X-tra adhesive coated microscopy slides, covered with kissol film and placed under pressure in a slide-press at 37°C for >48h. Tartrate resistant acid phosphatase (TRAP) and von Kossa staining were conducted as described previously and sections analysed using the bone histomorphometry software, HistoMorph (10).

*Raman Spectroscopy*

Raman spectra were acquired using an InVia™ Qontor™ Raman microscope (Renishaw, UK) equipped with a 633 nm HeNe laser and 1800 lines/mm grating (providing a spectral resolution of ~1 $cm^{-1}$) and a Master Renishaw Centrus 1UTR51 charged-coupled device (CCD). Prior to data acquisition, CCD and spectrometer slits were aligned using the auto-align function, and the laser spot was manually centred on the crosshairs using the camera. The system was intensity and wavenumber calibrated to the 520.7 $cm^{-1}$ peak of an internal silicon substrate. Single static point scans with an exposure time of 20s, 3 accumulations and 100% laser power (~31 mW at the sample) between 650–1750 $cm^{-1}$ within the "Fingerprint region" (11) were set using the WiRE 5.6 software (Renishaw, UK) and were acquired using a 20x air objective (NA of 0.4) yielding a diffraction limited spot size of ~965nm. A total of 75 spectra, presented as class means, were obtained from randomly selected locations from the tibial epiphysis in MMA-embedded sections (n=3/genotype/sex).

Cosmic ray artefacts were removed from individual spectra upon acquisition in WiRE 5.6, as previously described (12). Subsequent spectral preprocessing and analysis was performed in iRootLab (13). Spectra were wavelet denoised (Haar-wavelets, 6-point smoothing) prior to the fitting of a $5^{th}$-order polynomial for background fluorescence removal. Spectra were anchored to the x-axis using the rubber-band function and vector normalised. The second order derivative was calculated ahead of univariate spectral deconvolution in WiRE 5.6. The Raman peak corresponding to PMMA was identified at ~812 $cm^{-1}$ as assigned by (14). The peak positions corresponding to the major vibrations of ECM components, including proline (853 cm-1), hydroxyproline (876 cm-1), hydroxyapatite (960 cm-1), and B-type carbonate (1070 cm-1) were identified in second order derivative spectra. Peak areas were extracted by spectral deconvolution wherein a mixture of Lorentzian and Gaussian curves were fitted to class means spectra for specified Raman bands of interest.

*Backscattered SEM*

Right humeri were fixed in 4% paraformaldehyde, 2.5% glutaraldehyde (in 0.1 M sodium cacodylate) and stored in fixative at 4°C (15). Samples were bisected, dehydrated in an ethanol series, and embedded in epoxy resin (TAAB), and the surface polished using sequential grade diamond polishing strips (Agar Scientific). The block was mounted on a 25 mm stub, sputter coated with 10 nm Platinum using a Q150T ES coater (Quorum Technologies) and examined under back-scattered SEM (BSE-SEM) using a TESCAN Clara 2 field emission gun SEM equipped with a low energy annular backscattered detector and operating at 10 keV, at a working distance of 10 mm.

*Western blotting*

Femoral bone tissues were homogenised using a Rotor-Stator Homogenizer (IKA Ultra-Turrax T10) in radioimmunoprecipitation assay (RIPA) buffer containing protease inhibitors (Roche). Protein concentrations were measured by a BCA protein assay kit. The extracted proteins were equally loaded and separated using a 4-12% Bis-Tris protein gel and then transferred to a Polyvinylidene fluoride (PVDF) membrane with semi-dry transfer technique. The non-specific site of protein on the membrane was blocked with Intercept (TBS) Blocking Buffer (LI-COR Biosciences, Ltd.). Membranes were incubated sequentially in primary antibodies overnight at 4°C and secondary antibodies for 1 hour at room temperature (Table S2). β-actin conjugated with Horseradish Peroxidase (HRP) was incubated for 20 minutes at room temperature. The specific protein signals were detected by the ultra-sensitive enhanced chemiluminescence (ECL) substrates and imaged by the GeneGnome XRQ chemiluminescence imaging system (Syngene).

*Serum pyrophosphate ($PP_i$) and alkaline phosphatase (ALP) detection*

Plasma was collected by centrifugation at 2500 x*g* for 10 mins at 4°C following blood clotting for 30 mins at room temperature. $PP_i$ levels were measured using a colorimetric $PP_i$ assay kit (Abcam) according to the manufacturer's instructions. ALP was detected using a colorimetric ELISA kit (Novus Biologicals), as per the manufacturer's instructions. Absorbance was measured using a spectrophotometer (Biochrom EZ Read 2000) at 570 nm ($PP_i$) or 450 nm (ALP).

*Quantitative polymerase chain reaction (qPCR) and PCR array*

Left femora were cleaned of soft tissue, bone marrow removed by centrifugation and frozen at -80°C. Bones were thawed and homogenised using a Rotor-Stator Homogenizer (IKA Ultra-Turrax T10) in ice-cold Qiazol reagent. RNA was isolated and purified using a Qiagen RNeasy mini kit, according to the manufacturer's instructions.

RNA (2 µg) was reverse transcribed using a Qiagen QuantiTect RT kit, according to the manufacturer's instructions. The PCR was performed using cDNA (50ng), a Brilliant II Sybr green mastermix (Agilent), and run on an Agilent AriaMx PCR machine. Gene expression data for the qPCR were normalised to *β-actin* and *Atbp5* using the geometric mean and analysed using the ΔΔCt method (16). Primer sequences can be found in Table S3.

*Statistical analysis*

Analyses were performed using GraphPad Prism software (version 10.1.2). Normal distribution of data was assessed using the Shapiro-Wilk normality test. For comparison between genotypes, a one-way ANOVA followed by Tukey's post hoc analysis was used to test for significance. Results are presented as the mean ± standard error of the mean (SEM).

For the cortical bone analysis, R software (version 4.2.2) was used to generate line graphs and perform statistical analyses. Normal distribution of data was assessed using the Shapiro-Wilk normality test. For comparison between genotypes, a one-way ANOVA followed by Tukey's post hoc analysis was used to test for significance. Lines represent mean, with shading representing ± SEM. Colour heatmaps display the statistical significance at matched locations along the graph showing the genotype interaction and comparisons between genotypes. Red = $p<0.001$, green = $p<0.01$, yellow = $p<0.05$, grey = $p>0.05$ (not significant).
